# Evaluation of the 3D fractal dimension as a marker of structural brain complexity in multiple-acquisition MRI

**DOI:** 10.1101/124206

**Authors:** Stephan Krohn, Martijn Froeling, Alexander Leemans, Dirk Ostwald, Pablo Villoslada, Carsten Finke, Francisco J. Esteban

**Affiliations:** Department of Neurology, Charité Universitätsmedizin Berlin, Germany; Berlin School of Mind & Brain, Humboldt-Universität zu Berlin, Germany; Computational Cognitive Neuroscience Laboratory, Freie Universität Berlin, Germany; Department of Radiology, University Medical Center Utrecht, Netherlands; Image Sciences Institute, University Medical Center Utrecht, Netherlands; Center for Adaptive Rationality, Max-Planck Institute for Human Development, Berlin, Germany; Center of Neuroimmunology, Institut d’Investigacions Biomediques August Pi Sunyer (IDIBAPS), Barcelona, Spain; Systems Biology Unit, Department of Experimental Biology, Universidad de Jaén, Spain

**Keywords:** fractal analysis, structural brain complexity, MRI biomarker, structural similarity, imaging validation

## Abstract

Fractal analysis represents a promising new approach to structural neuroimaging data, yet systematic evaluation of the fractal dimension (FD) as a marker of structural brain complexity is scarce. Here we present in-depth methodological assessment of FD estimation in structural brain MRI. On the computational side, we show that spatial scale optimization can significantly improve FD estimation accuracy, as suggested by simulation studies with known FD values. For empirical evaluation, we analyzed two recent open-access neuroimaging data sets (MASSIVE and Midnight Scan Club), stratified by fundamental image characteristics including registration, sequence weighting, spatial resolution, segmentation procedures, tissue type, and image complexity. Deviation analyses showed high repeated-acquisition stability of the FD estimates across both data sets, with differential deviation susceptibility according to image characteristics. While less frequently studied in the literature, FD estimation in T2-weighted images yielded robust outcomes. Importantly, we observed a significant impact of image registration on absolute FD estimates. Applying different registration schemes, we found that unbalanced registration induced i) repeated-measurement deviation clusters around the registration target, ii) strong bidirectional correlations among image analysis groups, and iii) spurious associations between the FD and an index of structural similarity, and these effects were strongly attenuated by reregistration in both data sets. Indeed, differences in FD between scans did not simply track differences in structure per se, suggesting that structural complexity and structural similarity represent distinct aspects of structural brain MRI. In conclusion, scale optimization can improve FD estimation accuracy, and empirical FD estimates are reliable yet sensitive to image characteristics.

## 1 Introduction

Fractal analysis has attracted increasing interest from the neuroscience community as a versatile new tool for the analysis of structural brain data on a cellular as well as a macroscopic scale and in both health and disease (Di Ieva et al., 2014a, 2015, 2016). Fractal geometry, prominently developed by Benoît B. Mandelbrot (Mandelbrot, 1983), features the fundamental insight that real-world objects do not adhere to the smooth whole-integer dimensions of Euclidean geometry and are instead more adequately described by the fractal dimension (FD), which is not limited to integers and can be regarded as a measure of morphometric complexity (Mandelbrot, 1967; Di Ieva et al., 2016). While natural objects are constrained to finite physical scales and their self-similarity is rather statistical than compositional, the analysis of an object’s fractal properties has proven insightful in a variety of fields, from the inanimate (e.g. coastlines, clouds, lightning) and the cellular (e.g. protein surfaces, viral receptor molecules, cellular shapes) up to the realm of higher-order organisms (e.g. human bronchial and vascular ramifications) (Di Ieva et al., 2014a, 2015, 2016; Mandelbrot, 1983, 1967). In biomedical neuroimaging, fractal analysis has been applied in the anatomical description of cortical geometry (Kiselev et al., 2003; Im et al., 2006), and the fractal dimension has shown promise as a biomarker in the detection of early tissue alterations in multiple sclerosis (Esteban et al., 2007, 2009), brain abnormalities in infants with intrauterine growth restriction (Esteban et al., 2010), atherosclerotic white matter lesions (Takahashi et al., 2006), morphological changes in multiple system atrophy of the cerebellar type (Wu et al., 2010), angioarchitecture of cerebral arteriovenous malformations (Di Ieva et al., 2014b), the cortical features in Alzheimer’s disease (King et al., 2009, 2010; Ruiz de Miras et al., 2017), cerebral tumors (Iftekharuddin et al., 2009) as well as age-related brain atrophy (Madan and Kensinger, 2016) and age-induced structural changes in white matter tissue (Reishofer et al., 2018).

However, while fractal analysis is now being applied in both fundamental research and clinical investigations, there is a relative scarcity of literature on the methodological evaluation of the fractal dimension in structural brain MRI. On the computational side, one aspect that warrants further study regards the optimal range of spatial scales for empirical estimation, i.e. the regression intervals applied to the log-transformed data, specifically with respect to the commonly applied 3D box-counting procedure. In this context, we here propose a simple spatial optimization algorithm that automatically selects the optimal scale range for each individual estimation, and we present a series of simulation studies with known expected fractal dimensions to examine performance against non-optimized estimation.

Empirically, further examination is warranted with regard to the impact of fundamental image characteristics on the fractal dimension estimates, for instance regarding segmentation procedures, tissue type, image complexity, image registration, and spatial resolution. Moreover, it is important to assess the stability of the fractal dimension over multiple repeated acquisitions, since a reasonable test-retest reliability is an essential prerequisite for a biomarker’s diagnostic capacity. Furthermore, T1-weighted images (T1WI) have been the mainstay of neuroimaging studies implementing fractal analysis so that systematic evaluation regarding the utility of T2-weighted images (T2WI) in fractal analysis is comparatively scarce, even though the latter are essential to both fundamental neuroimaging research and clinical neuroradiological assessment.

To address these empirical questions, we analyzed structural MRI data from two independent openly available neuroimaging datasets (see below for details). On the one hand, this includes the recently published MASSIVE database (Multiple Acquisitions for Standardization of Structural Imaging Validation and Evaluation, cf. Froeling et al. 2017), featuring ten repeated T1WI and T2WI acquisitions over a short amount of time. We hypothesized that in such an acquisition procedure, it is reasonable to assume that there was essentially no change in the underlying structural brain complexity and that, therefore, the estimated fractal dimension values should show high stability across these short-interval measurements, allowing for detailed parameter-dependent analyses. While this data set is thus well-suited to examine the above questions, it also emanates from a single subject, potentially restricting the generality of our findings. Therefore, we extended our analyses to the recently presented Midnight Scan Club (MSC) data set (Gordon et al., 2017), featuring repeated short-interval acquisitions of T1WI and T2WI in 10 subjects. Our approach to the points raised above then rests on an image processing procedure differentiating between sequence weighting, spatial resolution, segmentation method, tissue type, and image complexity. As detailed below, this leads to a stratification of 32 distinct image analysis groups as a combination of image characteristics and processing parameters. We then apply fractality estimation with spatial scale optimization on the 3-dimensional input volumes obtained from image processing and implement a detailed and systematic analysis of the resulting fractal dimension estimates. The latter features a combination of random and systematic resampling methods, deviation detection, assessment of the sample distributions, similarity comparison, unsupervised machine learning techniques, correlation analyses, and parameter-dependent group comparisons. Based on these analyses, we assess 1) parameter-dependent repeated-sampling deviations, both within analysis groups and across the two data sets, 2) the impact of image registration on the fractal dimension estimates, 3) the within- and across-subject FD sample distributions, 4) the estimated optimal spatial scales across data sets, subjects, and processing parameters, 5) the relationship between the fractal dimension and structural similarity, and 6) the impact of image weighting, spatial resolution and processing parameters on the fractal dimension estimates.

## 2 Methods

### 2.1 Image acquisition and processing

Structural MRI in the MASSIVE data set were acquired on a clinical 3 T system (Philips Achieva). The data emanate from a healthy 25-year-old female subject scanned in five sessions occasions over an interval of 2 weeks. Ten T1WI and T2WI were collected, each reconstructed with 1 mm^3^ isotropic resolution, and data for both weightings were resampled to 2.5 mm^3^ isotropic resolution, resulting in the four image categories T1 high resolution, T1 low resolution, T2 high resolution, and T2 low resolution for further processing. Data were registered to a common space using a rigid registration algorithm (http://elastix.isi.uu.nl, see Klein et al. 2010; Shamonin et al. 2014) with the first T1 volume as the registration target. For additional details on the acquisition procedure, please refer to Froeling et al. 2017. The MASSIVE data set is openly available from www.massive-data.org.

Structural MRI in the MSC data set were obtained on a 3 T scanner (Siemens TRIO) across two separate days, with each session starting at midnight. Four T1 and four T2 scans with 0.8 mm^3^ isotropic resolution were acquired in each of the ten healthy subjects (5 female, 5 male; age range: 24-34 years). Additionally, subject #8 had one extra T1 scan, and subject #6 had five T1 scans and six T2 scans in total, which we included in our analyses wherever feasible. Similar to the above, data were resampled to a 2.5 mm^3^ isotropic resolution, and subject-wise rigid-body registration to the respective subject’s first T1 volume was carried out. For further details on the data set, see Gordon et al. 2017. The MSC data set is openly available from https://openneuro.org.

A standard FSL-based pipeline (Smith et al., 2004; Woolrich et al., 2009; Jenkinson et al., 2012) was used to preprocess the MR images for subsequent fractal analysis. Specifically, the brain extraction routine (BET) was applied to all individual 3D volumes with default fractional intensity threshold (Smith, 2002). The brain-extracted images entered the FAST routine for tissue segmentation into gray matter (GM), white matter (WM) and cerebrospinal fluid classes with default analysis parameters (Zhang et al., 2001). Intensity inhomogeneity was accommodated by iterative bias-field correction. We estimated partial volume maps for each of the three tissue classes, of which the GM and WM estimates entered the fractal analysis. For qualitative comparison, we also included a forced-decision binary classification (“hard” segmentation), in which voxels are labeled as 0 or 1 for a specific tissue class. Based on these segmentations, 3D image skeletons were estimated for each input volume. Image skeletons are the result of iterative reduction process that computes a minimum complexity version of the input image. We here apply a publicly available 3D parallel thinning algorithm to build the skeleton models of the respective input volume (Lee et al., 1994; Kerschnitzki et al., 2013). Intuitively, image skeletons aim at capturing the “essence” of an image and are thought to be more sensitive to pathological changes in some cases (Esteban et al., 2007, 2009, 2010; Jiménez et al., 2014), which is why we include them in the present study. For every input volume, we thus obtain eight resulting models. In summary, this amounts to a total of 32 analysis groups as a result of factorial combination, on which we base the taxonomy applied throughout the manuscript: image weighting (T1 vs. T2), spatial resolution (low vs. high), segmentation procedure (partial volume estimates (pve) vs. binary segmentation (bin)), tissue type (gray matter (GM) vs. white matter (WM)), and image complexity reduction (skeletonized vs. unskeletonized images, where the former is abbreviated by “Skel”). Figure 1 summarizes the analysis stratification (panel A) and provides an example of the processing results (panel B) as well as a 3D rendering of the corresponding skeleton models (panel C).

**Figure 1:**
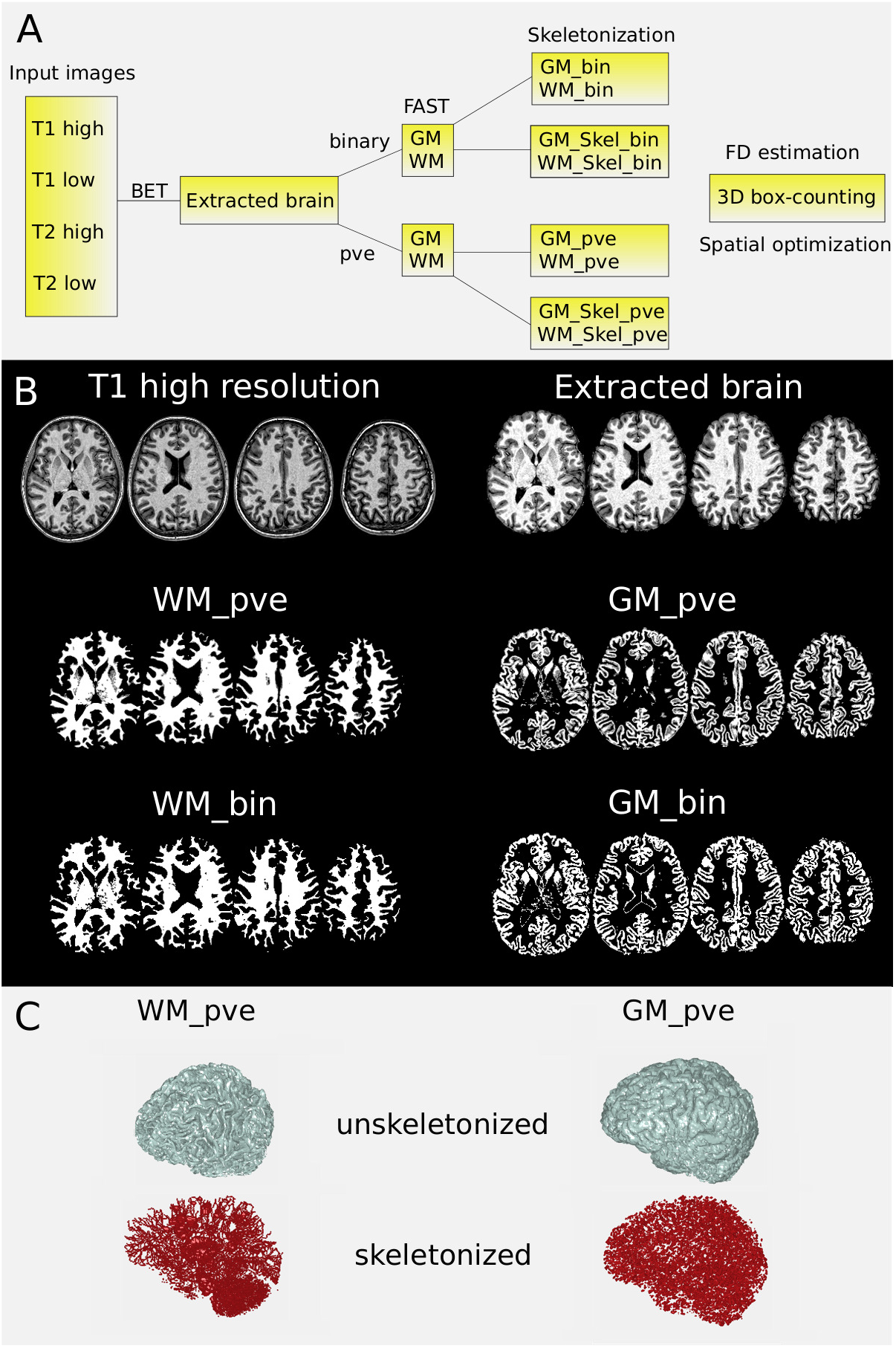
Analysis stratification and image processing. Panel A represents a schematic of the applied analysis stratification. An example of this procedure is visualized for the first volume of the T1 high resolution images in the MASSIVE data set (panel B). Note the absence of gray voxels in the binary forced-decision segmentations (bin) as compared to the partial volume estimates (pve). For each processed volume, image skeleton models were estimated, a 3D rendering of which is visualized for the WM_pve and GM_pve segmentations in panel C. The 3D volumes then entered the fractal dimension estimation. bin: binary segmentations; GM: gray matter; pve: partial volume estimates; Skel: skeleton model; WM: white matter.

### 2.2 Fractal estimation and spatial optimization

The volumes obtained from the above preprocessing provided the input for the estimation of the 3D fractal dimension. In the empirical sciences, the fractal dimension of an object *A* is commonly estimated by the box-counting dimension *D_b_* given by

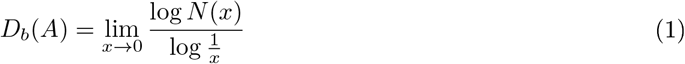

where *x* is the box edge length and *N* (*x*) the minimum number of boxes needed to cover the object under scrutiny (cf. Di Ieva et al. 2014a). Box-counting was applied here based on a function from the openly available calcFD toolbox (Madan and Kensinger, 2016). Due to the finite physical scales of natural objects, *D_b_*(*A*) is in practice calculated as the slope of the linear regression line over an interval of *x* in the log-log plot (see Gneiting et al. 2012 for detailed treatment of the ordinary least squares regression fit in box-counting). In terms of structural MR images, these intervals correspond to the range of voxel unit edge sizes over which the box-counting dimension is computed. In this context, consider a finite sequence *X_k_* of spatial scales defined as

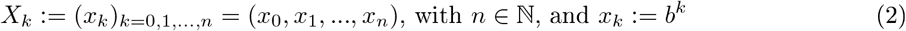

where *b* defines a scale base and *k* specifies the exponents to be tested.

For instance, we here define *b* = 2 and *k* = 0, 1, …, 8, yielding *X_k_* = (1, 2, 4, 8, 16, 32, 64, 128, 256). Nonetheless, the above raises the question over which particular range of *k* (i.e. which subsequence of *X_k_*) one should compute the box-counting regression in order to obtain the best fractal dimension estimate. One common solution is to simply define the *k*-range for the estimation and keep it fixed over repeated estimations. This, however, entails the danger of introducing inaccuracies as it disregards potential differences between subjects, scanning sessions, or processed input volumes. Another option is to base the definition on prior validation studies suggesting an optimal range of *k* for a particular image analysis group (see e.g. Esteban et al. 2009; Jiménez et al. 2014). Albeit an improvement, optimal spatial scales may depend on the scanning equipment, image processing, or estimation algorithm applied, and there is no principled reason to believe that the best regression intervals generalize uniformly from one population to another. As such, a more flexible and data-driven decision criterion may be desirable. We here apply a simple procedure to help alleviate this issue. Let *|X_k_|* denote the number of elements in the sequence of spatial scales resulting from eq. 2, and let *ω ≤ |X_k_|* indicate the upper bound on regression interval length, with *ω* = *|X_k_|* representing the case in which we allow estimation over all spatial scales in *X_k_* (i.e. here, *k* = 0, …, 8). However, we may also estimate the fractal dimension over a subsequence of spatial scales (e.g. *k* = 2, …, 5). Let *τ ≥* 2 denote the lower bound on the number of elements in this subsequence, i.e. the minimum length of the regression interval over which fractality estimation is carried out. The number of spatial scales of at least length *τ* and at most length *ω* is then given by the number of subsequences of *X_k_*, i.e.

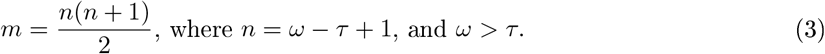

For a specified lower and upper bound on the regression interval, we thus obtain *m* possible *k*-ranges over which to carry out the estimation, yielding a set of *m* regression models. From this set, we may then choose the best-fitting model as suggested by the highest adjusted coefficient of determination 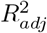, where standard adjustment (Fritz et al., 2012) is applied due to the varying cardinality of the different tested *k*-ranges. The slope estimator of the thus selected model is then chosen as the optimal fractal dimension estimate, in the sense of being the best guess in terms of approximating the true but unknown underlying dimension value based on the box-counting results.

In this context, empirical estimation faces the challenge that there is no obvious ground truth regarding the measure of interest. Specifically, as the true underlying fractal properties of the natural object under scrutiny are unknown, it is intrinsically difficult to judge estimation accuracy. As such, in order to examine the performance of the outlined procedure, we ran a series of simulation studies, in which we applied the estimation process to objects whose fractal dimension is known and can thus serve as a benchmark. Specifically, we created a series of 3D random Cantor sets whose expected fractal dimension is specified by the probability of retaining a particular subset during iterative removal (Falconer and Grimmett, 1992; Moisy, 2008). For each random Cantor set, we then estimate its fractal dimension over both the respective optimal spatial scales and over a randomly chosen non-optimal interval. Following initial parameter search in fixed benchmarking objects, we here apply *τ* = 4 (i.e. computing the regression over at least four contiguous spatial scales; *τ* = 3 yielded similar outcome) and *ω* = *|X_k_|* = 9 (i.e. allowing a maximum interval over all examined spatial scales), leading to *m* = 21 different models based on eq. 3. Figure 2 relates the corresponding simulation results: panel A displays the exemplary estimation of a non-fractal object (cube with expected *FD* = 3) and a fractal object (3D random Cantor set with expected *FD ≈* 2.7655). Compared to random non-optimal spatial scales, the outlined procedure improved estimation accuracy by several orders of magnitude (e.g. the arbitrary *k*-range estimates the Cantor set correctly to the first decimal, while the optimal *k*-range first deviates from the expected FD only in the fourth decimal place), even though 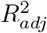 was very high in both cases. We then conducted a systematic simulation study, for which we created *n* = 100 distinct random Cantor sets for eight different retainment probabilities (from *p* = 0.6 to *p* = 0.95) yielding expected FD values in the range between 2 and 3 (with the aim of covering a biologically plausible range for fractal dimension estimates in brain MRI). Panel B displays the results of the subsequent fractality estimation over optimized and random non-optimal scales. Here, the proposed spatial optimization procedure produced improved estimation results in virtually all simulation iterations and for all expected fractal dimension values. In contrast, choosing a non-optimal spatial scale led to both pronounced over- and underestimation of the expected FD, and comparing estimation variance with Levene’s test suggested that optimized estimation precision was superior to non-optimal spatial scales at *p* < 0.001 for all retainment probabilities (right subpanel). In the following, we applied the same optimization procedure during fractal dimension estimation of the empirical data, and we explicitly analyze the estimated optimal *k*-ranges in the two data sets below.

**Figure 2:**
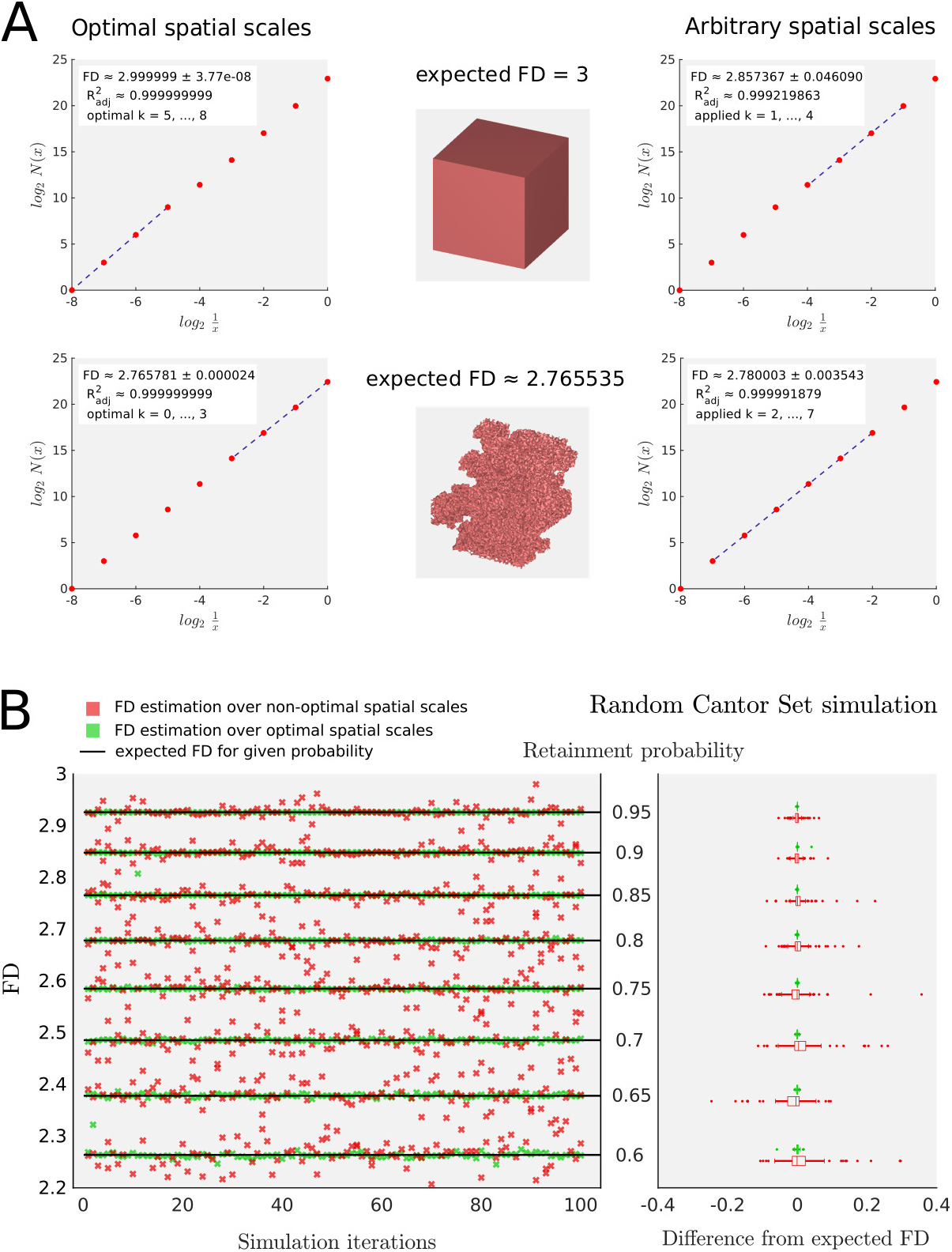
Fractal dimension estimation with spatial scale optimization. Panel A contrasts estimation results over optimal and arbitrary *k*-ranges for a non-fractal (cube) and a fractal object (3D random Cantor set) whose expected fractal dimension values are known. In both examples, optimization increases estimation accuracy by several orders of magnitude. Panel B displays the results of a random Cantor set simulation over varying retainment probabilities, yielding different expected fractal dimensions. Green crosses correspond to the outlined optimization procedure, while red crosses indicate estimation results over randomly chosen non-optimal spatial scales. The right subpanel relates the difference from the respective expected fractal dimension values over all estimations for the different retainment probabilities. Choosing a non-optimal spatial scale led to both pronounced over- and underestimation of the expected FD, and optimized estimation precision was superior to non-optimal spatial scales at *p <* 0.001 for all retainment probabilities.

### 2.3 Data analysis

#### 2.3.1 Deviation analysis

With the outlined processing stratification, we obtained a total of 320 fractal dimension estimates in the MASSIVE data set (10 subject scans x 32 analysis groups) and 1344 fractal dimension estimates in the MSC data set (42 subject scans x 32 analysis groups). In order to qualitatively assess the data within each analysis group, we first applied a combination of random and systematic resampling procedures. Specifically, we performed a bootstrapping procedure in order to randomly sample the mean and the 99 % normal approximation confidence interval (CI) of the FD over 2000 resampling iterations. Bootstrapping provided an objective way of qualitative data assessment in terms of the tightness of the confidence interval, which served as an indicator for the deviations within the analysis group, and the presence or absence of a skew in the clusters of the resampled means, indicative of important singular deviations in the raw estimates. Moreover, the bootstrapped CI was subsequently assessed as one of several criteria to identify meaningful deviations in the sampled FDs within each analysis group. We then applied a jackknife procedure, in which we systematically resampled the means by iteratively omitting each of the scans within the group in order to see if the variance changed significantly as assessed by Levene’s tests. We then made the explicit assumption that the FDs obtained within each analysis group were sampled from a true but unknown normal distribution. We fitted a Gaussian distribution to the sampled FDs and assessed the coherence to a corresponding theoretical distribution by means of a quantile-quantile plot. In order to examine whether the sampled data was reasonably assumed to follow a normal distribution, we furthermore computed the Shapiro-Wilk test (Shapiro and Wilk, 1965), applicable to assess composite normality for smaller sample sizes.

As an example, figure 3 visualizes these analysis steps for the exemplary group of binarized and skeletonized WM images in the T2 low resolution category (T2 low WM_Skel_bin) in the MASSIVE data set. The same analysis steps were applied to all 32 analysis groups in both data sets. In doing so, we sought to define a sensible criterion of when to “flag” an FD value due to a meaningful deviation within an analysis group. To this end, we compared various measures to find a balanced trade-off between detection and discrimination ability. First, we assessed whether a single FD value was inside or outside the bootstrapped confidence interval. As a second method, we assessed whether a particular value was within one or respectively two standard deviations (SD) of the sample mean. Third, we assessed whether the variances of the jackknife means significantly differed from one another by evaluating Levene’s test. Furthermore, we computed the Grubbs test to detect outliers within a given analysis group (Grubbs, 1969). The different methods were then assessed in terms of the original data and the effect that removing a flagged value had on the analysis in fig. 3. Specifically, we checked the flags against whether or not they occurred in groups in which the assumption of composite normality was first violated when considering all raw estimates, whether the removal of the flagged volume changed this, and if a deviation criterion would identify those analysis groups selectively. Based on the above points, the first method was deemed too conservative because the CI was tighter than even the one standard deviation interval of the sample mean and because it was sensitive to arbitrary choices regarding the type of computation (normal approximation vs. percentile-based, studentized or not, etc.). Systematic resampling nicely showed the qualitative effect that a single volume had on the overall mean and its variance but resulted in limited sensitivity in multivariate testing, despite increased accuracy in case of non-normality. Jack-knife resampling was thus considered too liberal for our purposes given the cases of deviation-induced non-adherence to composite normality. When the 1 SD interval around the sample mean was considered, volumes were more selectively flagged. However, this criterion does not account for the range of the data scatter, which was generally very small within analysis groups. See for example fig. 3, where the data were sampled in the subdecimal scatter range of well under 0.03. As a result, scanning sessions were flagged with relatively low selectivity, which was alleviated by choosing a 2 SD interval around the sample mean. Even more selective, the Grubbs test procedure closely flagged non-adherence to composite normality, which was generally reversed after removal of the flag. Therefore this method was deemed the most appropriate criterion with the more conservative 2 SD method as a cross check. For an exemplary identification of a flag see fig. 4 relating the results for T1 high GM_pve images in the MASSIVE data set. Here, Grubbs testing flags the FD that corresponds to the first scanning session (note that the more conservative 2 SD criterion equivalently identifies this flag). Systematic resampling shows that omitting the flagged value causes an upward shift of the mean and reduces its variance but this does not reach significance level in multivariate testing. The flagged FD causes the assumption of composite normality to be invalid although the remaining samples tightly follow the reference for normality. Omitting the flag restores normality and clearly "tightens" the distribution (cf. panel D), while non-parametric distribution comparison was insignificant. Based on the results of the deviation analysis within each analysis group, we then examined the occurrence of flagged volumes by subjects, scanning session, image weighting, and processing parameters across the MASSIVE and the MSC data sets (see sec. 3.1).

**Figure 3:**
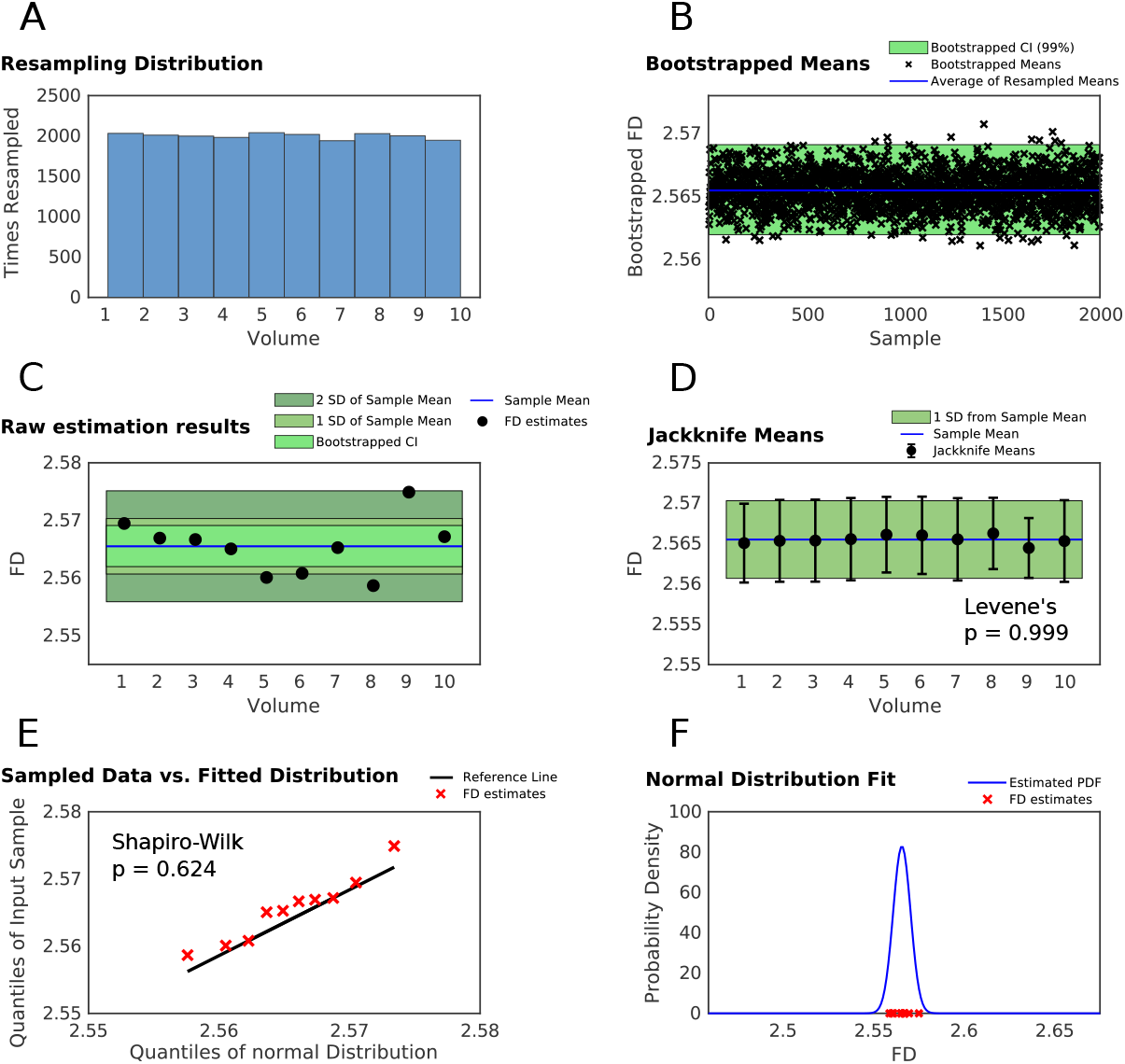
Main steps of within-group deviation analysis. The figure displays the deviation analysis for the exemplary analysis group of low-resolution T2 WM partial volume estimates in the MASSIVE data set. Panel A shows a near-uniform resampling distribution for bootstrapping, indicating the absence of *a priori* weights. Panel B displays the bootstrapped mean fractal dimensions as well as the resulting 99 % confidence interval and average over all bootstrapped means. Panel C plots the raw estimates for the ten scans in the data set and their sample mean, together with the bootstrapped confidence interval and the intervals spanning one and two standard deviations, respectively. Panel D represents the jackknife means, i.e. systematic resampling, where each of the ten raw was iteratively omitted to compute the mean over the remaining nine samples. Levene’s test to see if the variances of the thus obtained means significantly differed from one another was insignificant. Panel E shows a quantile-quantile plot for the original data vs. a fitted normal distribution, where a theoretical Gaussian would precisely follow the reference line. The values of the current analysis group reasonably adhere to this reference, and the test decision suggested that assuming composite normality was acceptable. Panel F shows the corresponding estimated normal distribution together with the cluster of the sampled FDs. The same procedure was applied to all 32 analysis groups in both the MASSIVE and the MSC data sets. CI: confidence interval; FD: fractal dimension; PDF: probability density function; SD: standard deviation.

**Figure 4:**
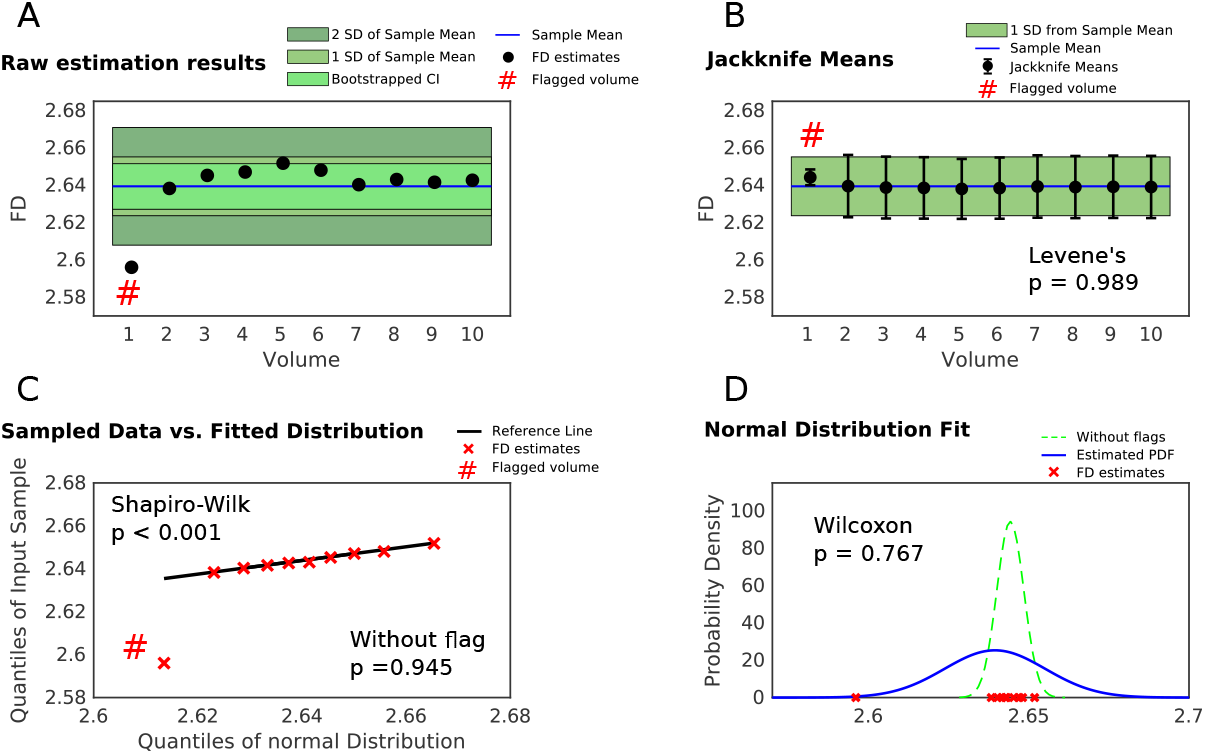
Exemplary identification of a within-group deviation. The data presented here belongs to the high-resolution T1 GM_pve images in the MASSIVE data set. If the fractal dimension of an image was identified to deviate from the remaining analysis group according to the chosen deviation criterion, the corresponding volume was flagged (indicated here by #). In this case, the FD value belonging to the first scan was flagged, and its deviation from the remaining samples is visible from panel A. Note that in panel B, the variance of the jackknife mean without this flagged volume is notably smaller, although this did not reach significance level in multivariate variance comparison. Panel C shows the corresponding quantile-quantile plot. Although the flagged FD only deviates by about 0.05 from the other FD estimates, normality assessment suggests that assuming an underlying Gaussian distribution is not recommendable. Clearly, however, the remaining samples tightly follow the normality reference and discarding the flagged FD indeed restores the acceptance of composite normality. Furthermore, non-parametric comparison between the fitted distributions with and without the flagged volume yielded insignificant results, exemplified here in panel D. CI: confidence interval; PDF: probability density function; SD: standard deviation.

#### 2.3.2 Impact of image registration

Based on the above analysis, we tested the effect of image registration and the ensuing interpolation on the fractal analysis results. In the MASSIVE data set, images were originally registered to the first T1 volume, and thus not all images were subject to the same transformations. To assess the impact of registration, we therefore reregistered all images to the mean of the FLAIR images, also included in the MASSIVE data set but independent of the presented analyses, and extended our analyses to the thus reregistered data. For further examination, we moreover reregistered the MSC data using FSL’s MNI152 structural template. We then compared the mean FDs in the 32 analysis groups between the respective first volume registration and the reregistered data non-parametrically by a series of Wilcoxon rank sum tests, with Bonferroni-Holm correction for multiple comparisons. Effect sizes for these comparisons are calculated based on the *z* value of the test statistic as 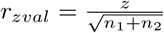, where *n*_1_ and *n*_2_ are the compared sample sizes, i.e. number of scans for the two respective registrations, cf. Fritz et al. 2012. Moreover, we computed correlation matrices to examine if there were associations between the 32 image analysis groups and whether image registration had an effect on potential associations.

#### 2.3.3 Fractal dimension and structural similarity

Furthermore, we sought to investigate the relationship between structural complexity and structural similarity. The motivation behind this was to examine if differences in fractal dimension essentially just track differences in structure, i.e. if two MRI volumes differ little in their fractal dimensionality simply if they are similar to one another. In this context, we computed the Structural Similarity Index (SSIM) between two given 3D volumes and related it to the difference of their respective fractal dimensions. The SSIM is a well-known reference metric of structural similarity between two images based on luminance, contrast and structure, and is commonly applied in signal processing and image quality assessment (Wang et al., 2004). The SSIM is bounded by [−1, 1], with *SSIM* (*x, y*) = 1 if the two images *x* and *y* to be compared are identical. Furthermore, the SSIM exhibits symmetry, such that *SSIM* (*x, y*) = *SSIM* (*y, x*) holds for any two images *x* and *y* (Wang et al., 2004; Østergaard et al., 2011; Brunet et al., 2012). We here computed the SSIM in every possible pair-wise comparison of two volumes within an analysis group (i.e. volume 1 vs. 2, volume 1 vs. 3, etc.) in both the MASSIVE and the MSC data set. The number of total unique comparisons between any two out of *n* input volumes is given by the binomial coefficient 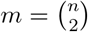, and thus we compute

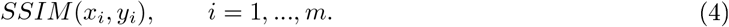

For each of these comparisons, we calculate the difference of the corresponding fractal dimension values of volume *x_i_* and *y_i_*, i.e.

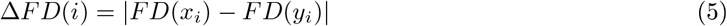

where we take the absolute difference to match the symmetry of the SSIM. In the MASSIVE data set, there are *n* = 10 repeated scans of a single subject. For each of the 32 analysis groups, we thus obtain *m* = 45 ∆*FD/SSIM* pairs, each belonging to one particular comparison of two 3D volumes. In the MSC data set, there are *n* = 4 repeated scans in each of the 10 subjects, yielding *m* = 6 between-volume comparisons in each analysis group. While the within-subject comparisons were thus considerably more limited, the MSC data set allowed us to extend the above question to across-subject analyses. To this end, we computed all possible session-wise comparisons between subject scans (i.e. session 1 subject 1 vs. session 1 subject 2, …, session 4 subject 9 vs. session 4 subject 10), yielding 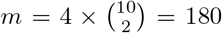 comparisons for each of the 32 analysis groups.

As plotting ∆*FD* over *SSIM* was suggestive of data clusters in some cases, we carried out a group-wise *k − means* clustering analysis. To this end, *k* was chosen agnostically based on range-constrained silhouette optimization (see appendix for an example and further details), yielding *k* = 2 for most analysis groups, followed by *k* = 3 in some instances. The clustering algorithm was run on the corresponding ∆*FD/SSIM* pairs with ten replicates to avoid convergence on non-global minima due to random initial conditions. Clustering quality was generally very good across the data sets as indicated by high average silhouette values and reasonably balanced cluster sizes. We furthermore examined whether there were significant associations between ∆*FD* and *SSIM* by means of non-parametric Kendall’s *τ* correlation, and performed a linear regression for all significant dependencies. In order to test if differences in fractal dimension induced by varying interpolation (see above) were related to structural similarity, and if the relationship between ∆*FD* and *SSIM* was altered due to different image registration, we conducted the above analysis in both the first volume and the reregistered data sets with identical optimization settings and compared ∆*FD*, *SSIM*, and *k − means* clustering results between the different registrations.

#### 2.3.4 Fractal dimension by image characteristics

Finally, we assessed differences of the fractal estimates across analysis groups as a function of image characteristics and analysis parameters. To this end, we compared the corresponding mean fractal dimensions by computing an analysis of variance (ANOVA), which invariably yielded significant differences in FDs across groups, and applied a post-hoc Tukey-Kramer test (Hayter, 1984) to investigate significant FD differences between analysis groups in pair-wise parameter-dependent comparisons. For all statistical tests employed in the present work, we defined a minimum significance level of *α* = 0.05.

Image processing was implemented with a set of Unix shell scripts. Skeletonization, spatial optimization studies, fractality estimation, and data analysis were carried out based on custom-written Matlab code (The MathWorks, Inc., Natick, MA, United States). For the interested reader wishing to retrace our analyses, all files are available from the Open Science Framework (http://osf.io/3mtqx).

### 3 Results

#### 3.1 Deviation analysis

The procedure detailed in sec. 2.3.1 was applied to all 32 analysis groups across the MASSIVE and MSC data sets, the result of which is shown in Figure 5. Generally, the overall robustness of the FD against repeated-sampling deviations was very high across both data sets, with over 95 % unflagged volumes. For the detected flags, our analyses uniquely identified a single scanning session that was responsible for the majority of deviations in both the MASSIVE and the MSC data sets in original registration, in this case volume 1 (figure 5, panels A and C). As the first T1 volume served as the respective subject-wise registration target, this finding motivated further examination in the reregistered data sets (cf. sec. 2.3.2). Interestingly, reregistration consistently abolished the clustering of deviations in the first volume in both the MASSIVE and the MSC data (panel B and D, respectively). Furthermore, reregistration further reduced the absolute number of deviations in both data sets by around 1.5-2 %. Despite this general reduction, reregistration also induced a few previously absent deviations in both data sets (e.g. volume 6 in the MASSIVE data; subject 7, volume 4, in the MSC data). In terms of image parameters, high resolution images were more susceptible to the effect of registration (with a slight predilection for T1WI), and skeleton models were more prone to deviations than unskeletonized images, while deviations were rather balanced between segmentation procedure and tissue type.

**Figure 5:**
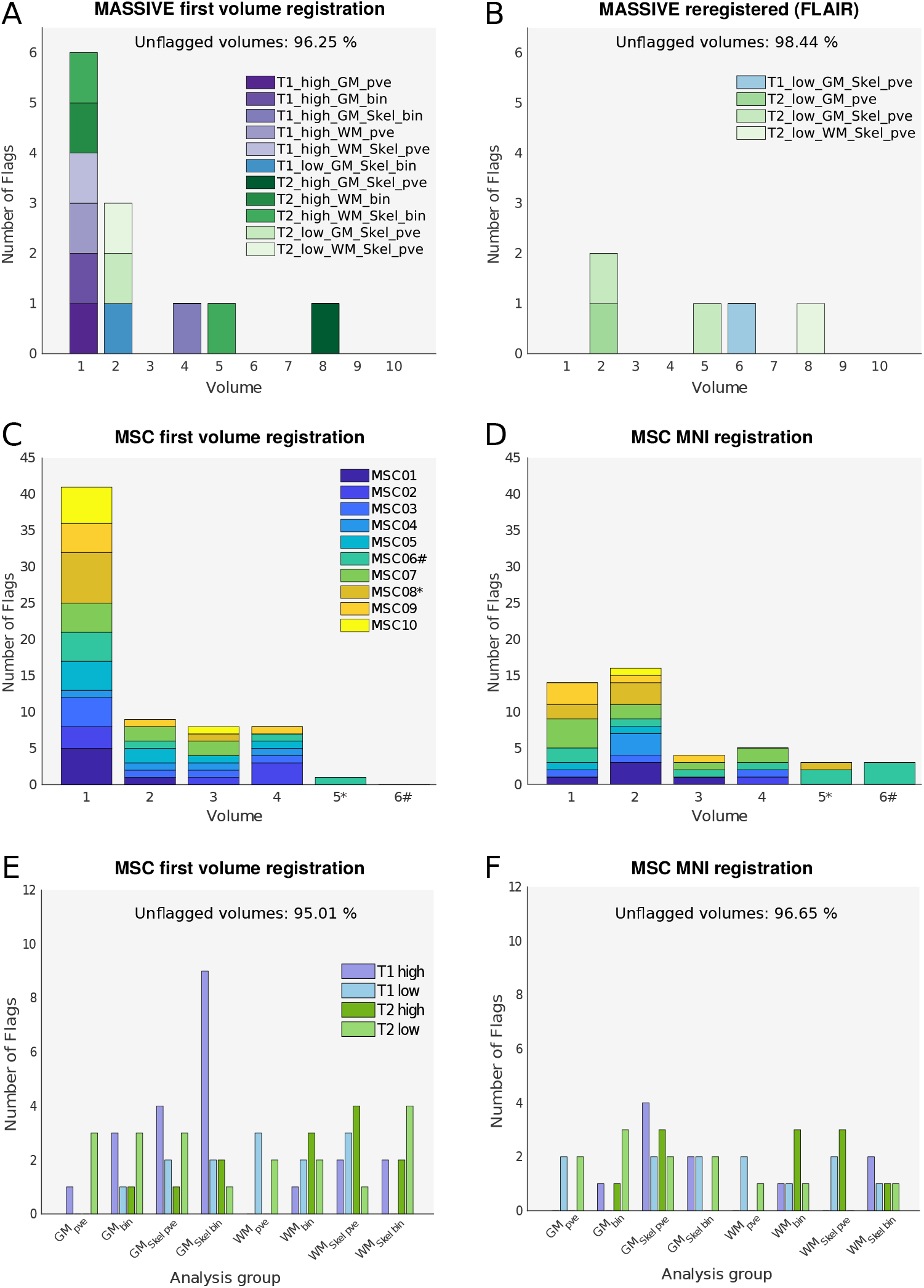
Deviation analysis across the MASSIVE and MSC data sets. Panels A and B depict sampling deviations by volume and analysis group in the MASSIVE data set in the original first volume registration and after reregistration to the mean FLAIR images. Panels C and D relate the results by volumes and subjects in the Midnight Scan Club (MSC) data in first volume and MNI registration. Note that only subjects 8 and 6 underwent acquisition runs 5 and 6, respectively (indicated by * and #), while all other subjects had four acquisition runs. Panels E and F resolve the MSC deviations by analysis groups in the two registrations. The original registration resulted in a deviation cluster around the registration target in both the MASSIVE and the MSC data. This effect was abolished by reregistration in both data sets. High-resolution images were more susceptible to the registration effect, and skeleton models were more prone to deviations than unskeletonized images. GM: gray matter; bin: binary tissue segmentation; pve: partial volume estimates; Skel: skeleton model; WM: white matter.

#### 3.2 Impact of image registration on fractal dimension profile

##### Absolute fractal dimension estimates

For further characterization of registration effects, we compared the fractal dimension profiles across all analysis groups between the two respective registrations for both data sets. As summarized in table I, image registration had a significant impact on the mean fractal dimension estimates for most analysis groups in T2WI for the MASSIVE data set, while the comparisons in T1WI were less often significant. For the MSC data set, all comparisons in the high resolution category for both T1WI and T2WI yielded significant results, while differences were less pronounced for low resolution volumes, especially in T1WI. Notably, in both registrations and both data sets, standard deviations for skeleton models across most analysis groups were up to one order of magnitude higher as compared to their unskeletonized counterparts (e.g. T1 low-resolution WM estimates). Moreover, data scatter was generally higher in the MSC data (across-subject means) as compared to the MASSIVE data (within-subject means). Regarding the direction of the effects, all significant registration-induced changes of the skeleton models in the MAS-SIVE data resulted in a decreased mean fractal dimension, i.e. reregistration uniformly reduced FD values in image skeletons. In contrast, the opposite pattern occurred in all but one of the unskeletonized image groups, with reregistration yielding higher mean fractal dimension estimates. Across the MSC data set, on the other hand, reregistration invariably resulted in decreased FD estimates for both T1 and T2 high resolution volumes, while mean FDs of low resolution images were generally increased. While registration-induced changes were thus quite consistent within each data set, the absolute mean values and the direction of registration-induced changes did not generalize from one data set to another.

**Table I:**
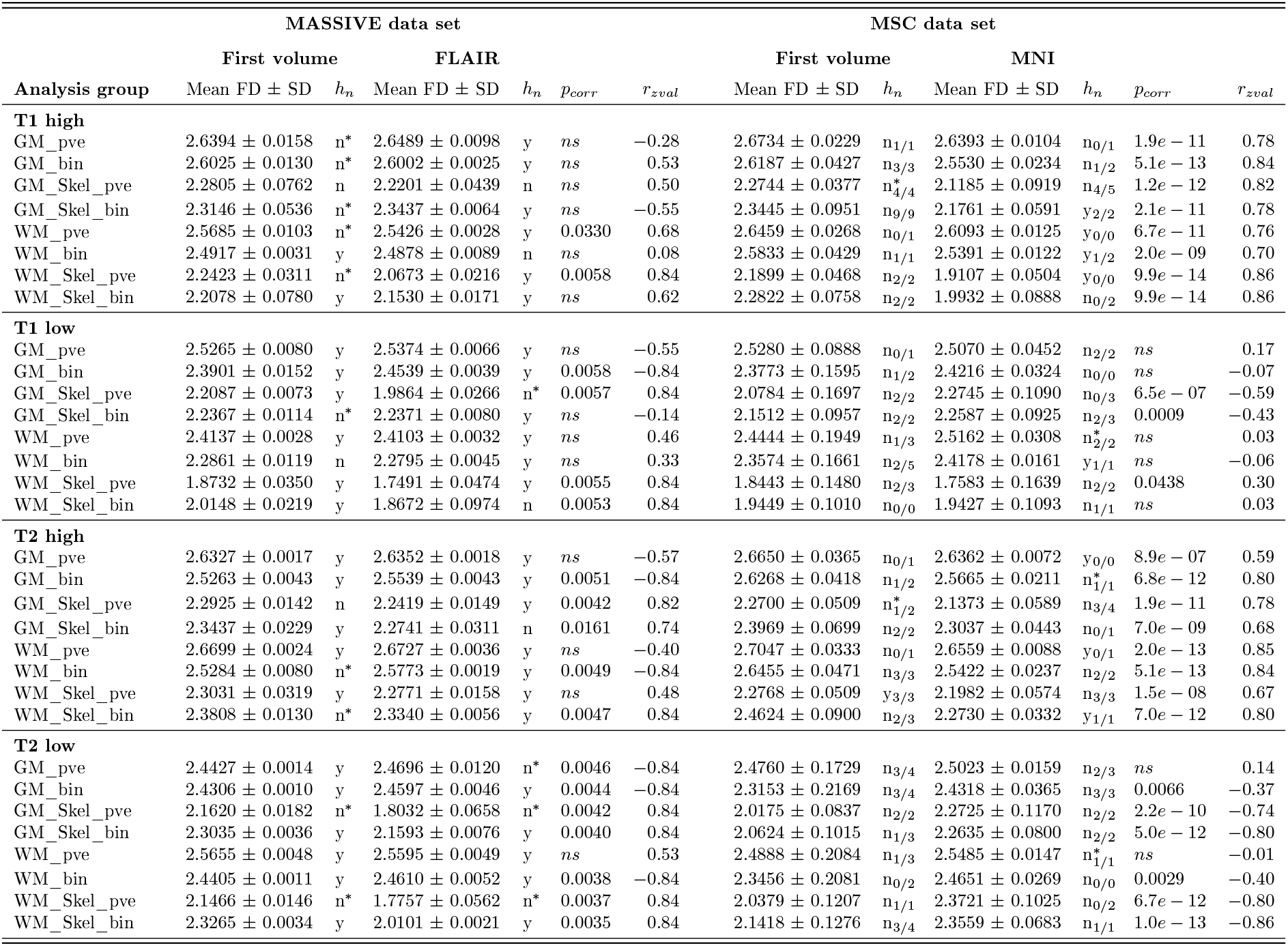
Impact of image registration on fractal dimension profile. The table summarizes the mean fractal dimension values by image group for the first volume registration and the reregistered data in both the MASSIVE and the Midnight Scan Club (MSC) data sets. Assessment of within-group composite normality (*h_n_*) is indicated by ‘y’ (yes) and ‘n’ (no). Asterisks indicate those groups in which composite normality was first violated but restored after removal of a within-group deviation, cf. sec. 3.1. Mean fractal dimensions between registrations were compared non-parametrically by Wilcoxon signed rank tests with Bonferroni-Holm-adjustment for multiple comparisons. Effect sizes are calculated based on the *z* value of the test statistic as *r_zval_* (cf. sec. 2.3.2). bin: binary segmentation; FD: fractal dimension; GM: gray matter; ns: not significant; p_*corr*_: adjusted p-value; pve: partial volume estimates; SD: standard deviation; Skel: skeleton model; WM: white matter.

##### Sample distributions

We furthermore assessed the sample distributions of the repeated-acquisition fractal dimension estimates in response to image registration across the two data sets. Specifically, table I summarizes the outcomes of composite normality assessment (*h_n_*) both within-subject (MASSIVE and MSC data) and across-subject samples (MSC data). Here, asterisks indicate the conversion cases, where composite normality was first refuted but accepted upon removal of the within-group deviations as identified by the deviation analysis from sec. 2.3.1 (cf. figs. 4 and 5). For the MSC data, the test decision refers to the sample across all subject volumes, with subscripts indicating how many within-subject normality assumptions were refuted without and respectively with these flagged volumes (a maximum of 10 for each analysis group based on the 10 subjects). As a general result, the normality assumption in within-subject measurements was more often refuted in first volume as compared to reregistration, although this reached significance level only for the MSC data (MASSIVE: 40.6% in first volume registration vs. 25% in FLAIR registration, *χ*^2^ = 1.1, *n* = 32, *p* = 0.29; MSC: 25.3% in first volume registration, 17.2% in MNI registration, *χ*^2^ = 5.8,, *n* = 320, *p* = 0.01). Furthermore, the repeated-sampling deviations constituted a main reason for a priori rejection of composite normality in within-subject sampling: in the MASSIVE data set, 10/13 normality rejections were restored by omitting deviations in first volume registration, and 4/8 in the reregistered data set. Conversion rates were 28.4% in first volume registration and 29.1% in MNI registration for the MSC data set. Considering the conversion cases, a total of 28/32 analysis groups adhered to composite normality in the reregistered MASSIVE data (87.5 %), with similar results for the within-subject distributions in the MNI-registered MSC data set (281/320 within-subject measurements, 87.8 %). While assuming an underlying normal distribution for within-subject sampling was hence acceptable for most analysis groups across both data sets, this did not transfer to the across-subject distributions in the MSC data set. Here, normality was refuted in the vast majority of analysis groups in first volume registration, and this was virtually unaltered by omitting within-subject deviations. MNI-registration yielded adherence to composite normality in 25 % of the analysis groups, without any obvious distribution across image categories, and this was again practically unaffected by within-subject deviations. Closer examination of the sample distributions suggested that reregistration had a discernible regularization effect on the across-subject distributions in some analysis groups, but not in others, as exemplified in fig. 6 for high-resolution gray matter partial volume estimates in T1WI and T2WI.

**Figure 6:**
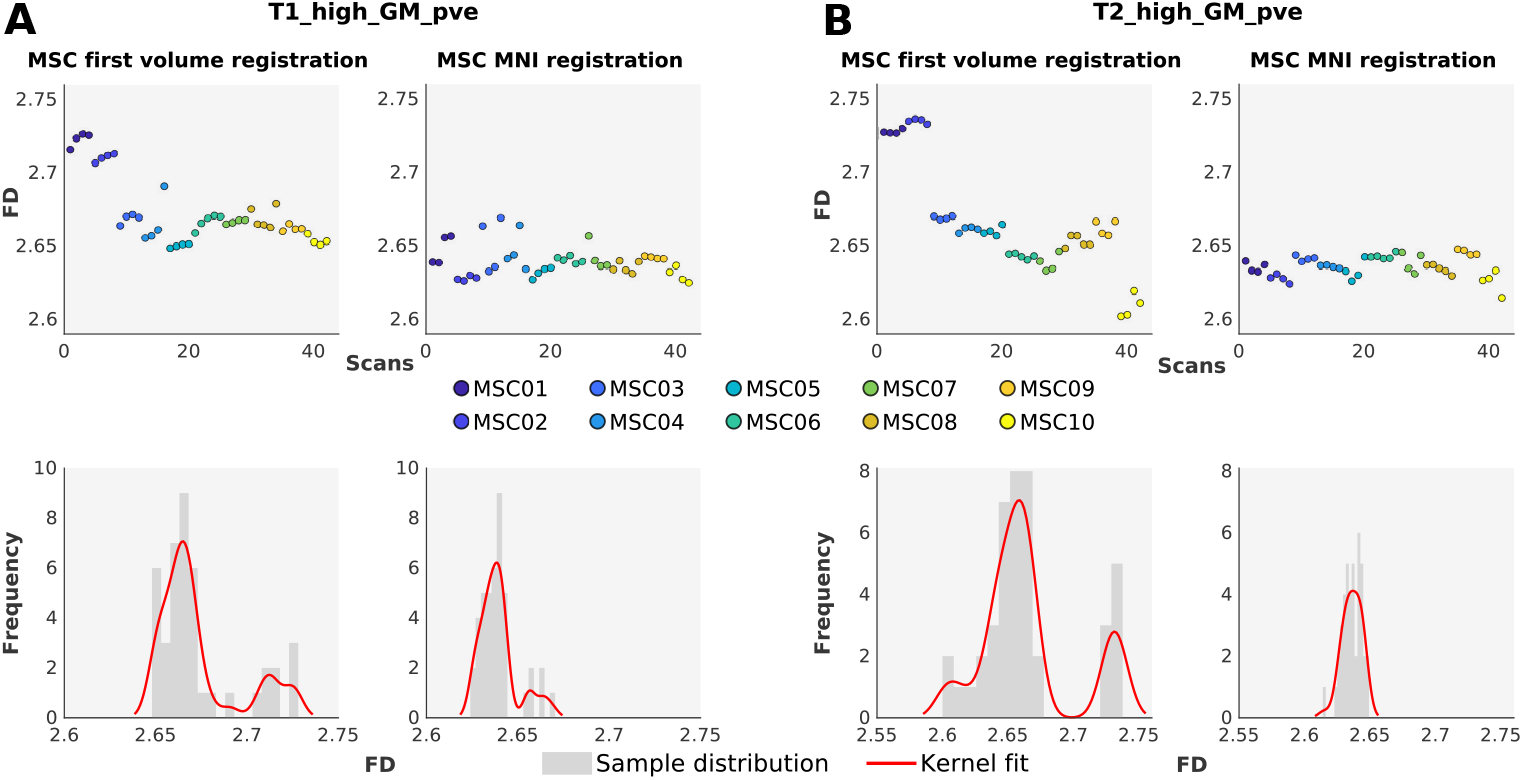
Exemplary across-subject distributions of fractal dimension estimates in the MSC data set. The figure reports the raw fractal dimension estimates by subjects, and the sample distributions with kernel density estimations across image registrations for the exemplary analysis groups of high-resolution gray matter partial volume estimates in T1WI (panel A) and T2WI (panel B), respectively. While MNI registration of the T2WI resulted in a regularization of the across-subject sample (and composite normality was acceptable), this was not the case for T1WI. GM: gray matter; pve: partial volume estimates.

##### Across-group associations

Based on the complex impact of image registration on the fractal dimension estimates in both data sets, we furthermore investigated whether there were any between-group associations across the 32 analysis groups and whether image registration had an impact on these associations. Figure 7 reports the corresponding results for the MSC data set (results for the MASSIVE data set were similar but limited to ten estimates in each group and only reflective of within-subject associations). First volume registration featured a large number of systematic, strong, bidirectional, and highly significant between-group correlations (panels A and C), reflected in a "checkerboard" pattern of the correlation matrix. Interestingly, reregistration to MNI space resulted in a pronounced overall across-group decorrelation, reducing both the strength and the amount of associations between image analysis groups (panels B and D), while an across-group association cluster was seen for some analysis groups in the T2 low-resolution category.

**Figure 7:**
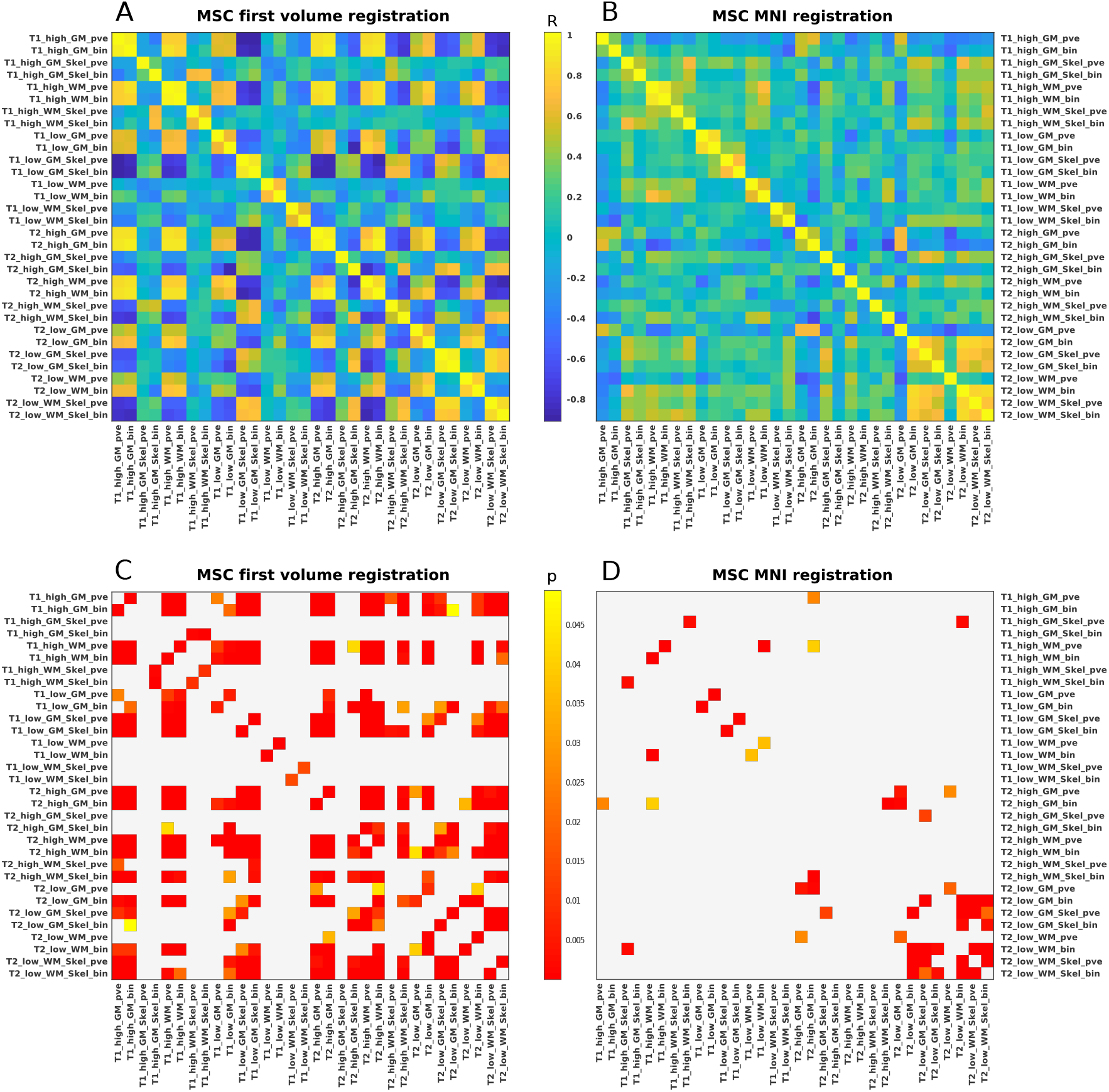
Across-group correlations in MSC data set. Panels A and B depict the correlation coefficients across the 32 image analysis groups in the Midnight Scan Club (MSC) data set in first volume and MNI registration, respectively. Panels C and D show the corresponding *p*-values below significance threshold after Bonferroni-Holm adjustment. While first volume registration induced strong systematic correlations between analysis groups, both the amount and the strength of these associations were markedly attenuated by reregistration. bin: binary tissue segmentation; pve: partial volume estimates; Skel: skeleton model.

#### 3.3 Optimal *k*-ranges

Building on the above results, we analyzed the optimization results across the two data sets in terms of analysis parameters and image registration. Specifically, for each individual fractality estimation, we tracked which spatial scale interval (i.e. which range of *k* in eq. 2) was selected as the optimal range for that particular estimation according to the procedure in sec. 2.2. Based on eq. 3, there were *m* = 21 distinct spatial scale intervals, ranging from *k* = 0, …, 3 to *k* = 0, …, 8. Figure 8 visualizes the frequency of the optimal *k*-ranges as estimated from the data. Panels A and B display the optimization results across analysis groups for the MASSIVE data set in first volume and FLAIR registration, respectively. As a general result, optimal *k*-ranges were highly selective in that they 1) displayed a clear preference for a subset of all possible spatial scales (i.e. were far from a uniform distribution), 2) differed markedly over the various analysis groups, and 3) showed a systematic tendency towards lower-cardinality over higher-cardinality scale intervals. Furthermore, the *k*-ranges in figure 8 are ordered from left to right by interval length and lower to higher *k*-values within each of these groups (i.e. from *k* = 0, …, 3 to *k* = 5, …, 8 for a cardinality of 4, from *k* = 0, …, 4 to *k* = 4, …, 8 for a cardinality of 5, and so on). From this it becomes apparent that optimal spatial scales showed a further tendency towards lower *k*-values (i.e. smaller box sizes) for a given interval length. For instance, considering a cardinality of 4, all estimations in the reregistered data set yielded optimal scales from *k* = 0, …, 3 to *k* = 3, …, 6, while the higher box edge sizes of *k* = 4, …, 7 and *k* = 5, …, 8 were never selected as optimal (panel B). Interestingly, scale selectivity in the MASSIVE data was even further increased by reregistration to the FLAIR images (in panel B, eleven *k*-ranges contained all optimization results, while the remaining ten were never chosen as the optimal spatial scales). Optimization outcome furthermore differed by image analysis groups. While there was no obvious distribution of optimal scales by weighting, resolution, segmentation procedure, or tissue type, a discernible pattern emerged as a function of skeletonization, on which we thus focus the visual comparison (with unskeletonized volumes in colder colors, and skeleton models in warmer tones). Optimal scales for image skeletons were systematically shifted to the right of unskeletonized images, i.e. intervals for skeleton models were generally of the same length but over higher *k*-values. Furthermore, we examined how consistently a particular *k*-range was selected in repeated estimations within the same image analysis group. To this end, we track how many volumes in each analysis group yielded the same optimal scale, regardless of the particular *k*-range. Panel C visualizes this scale dispersion for the MASSIVE data set. For some analysis groups in first volume registration, estimation yielded the same optimal scales for all ten input volumes, while there were nearly as many cases in which a *k*-range was only chosen once in a particular analysis group. Interestingly, reregistration shifted this distribution to the right, i.e. more analysis groups now consistently yielded the same optimization outcome over all ten input volumes. The same analyses were carried out over the MSC data set, summarized in panels D-F. Results closely mirrored the above findings in the MASSIVE data. Optimal *k*-ranges showed highly similar convergence on lower-cardinality intervals as well as lower *k*-values for a given interval length, with high consistency across subjects. Moreover, the same distribution of skeleton models and unskeletonized images was observed, and this pattern as well as scale selectivity was equivalently augmented by reregistration (panel E). Furthermore, the scale dispersion distribution in panel F was also right-shifted in the reregistered data set, indicating increased optimization consistency. This effect, however, was more pronounced in some subjects than in others, and absolute counts differed moderately among subjects, suggesting that despite high qualitative consistency, there was also some between-subject variability in the numerical frequency of individual optimization results.

**Figure 8:**
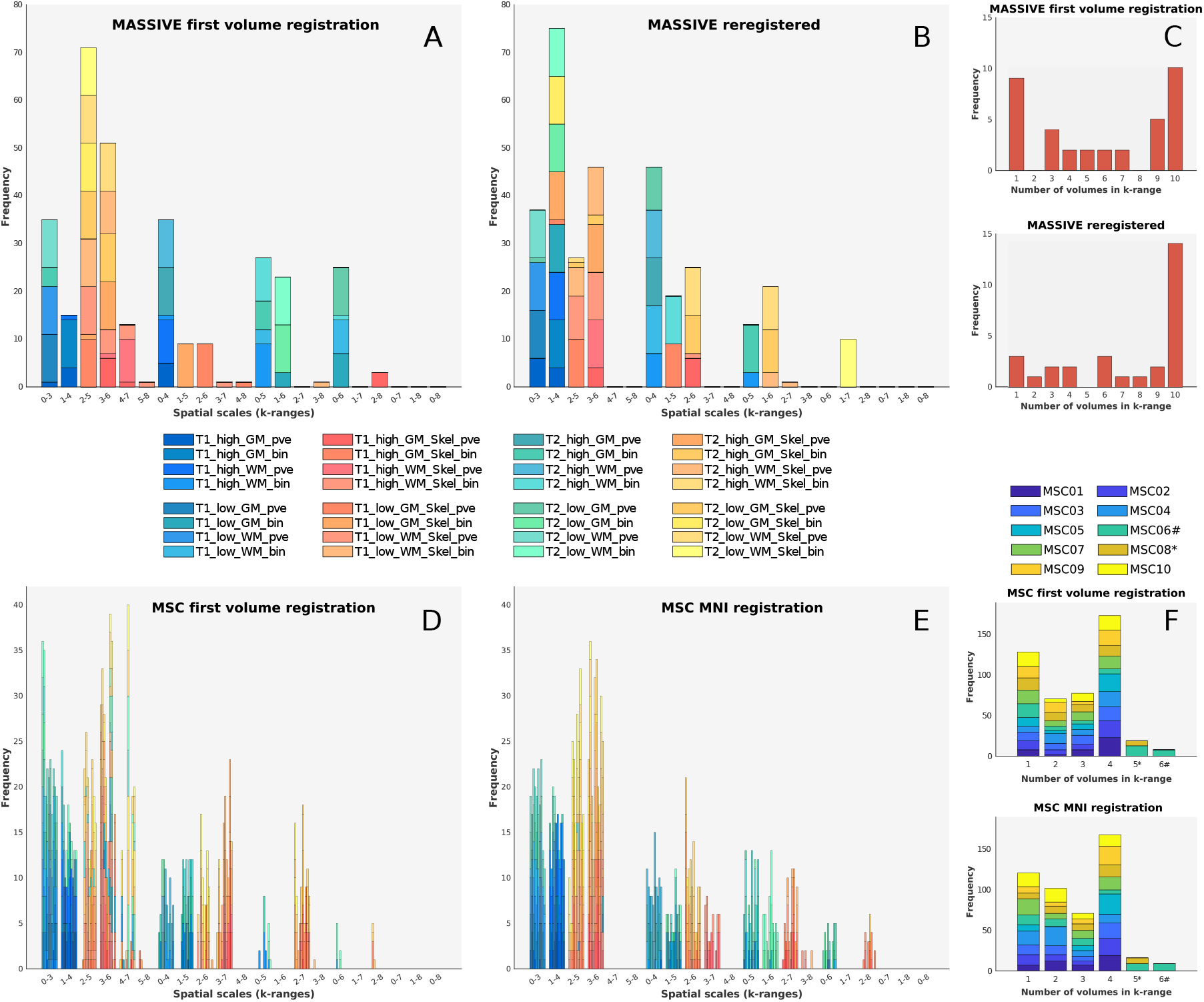
Optimal *k*-ranges in MASSIVE and MSC data sets. Panels A and B display the optimal spatial scales across all fractal dimension estimations in the MASSIVE data set for first volume and FLAIR registration. Panel C quantifies how many of the ten volumes in each image analysis group yielded the same respective optimal *k*-ranges as a measure of scale dispersion. Reregistration shifted this distribution to the right, reflecting increased consistency of repeated optimization results. Panels D and F show the absolute frequencies of optimal spatial scales in the MSC data set for first volume and MNI registration (single bars represent subjects and stacks represent image analysis groups for each subject). There was notable similarity to the MASSIVE data in scale selectivity and distribution by image analysis groups, especially regarding skeleton models vs. unskeletonized images. Panel F represents the consistency distribution over subjects in the MSC data. Note that only subjects 8 and 6 underwent acquisition runs 5 and 6, respectively (indicated by * and #), while all other subjects had four acquisition runs, and thus 4 represents the maximum repeated-optimization consistency for those subjects. GM: gray matter; bin: binary tissue segmentation; pve: partial volume estimates; Skel: skeleton model; WM: white matter.

#### 3.4 Fractal dimension and structural similarity

The procedure in sec. 2.3.3 revealed an interesting relationship between the fractal dimension and structural similarity. Generally, *SSIM* values were found in the range of 0.7 and 1 for both data sets, indicating a high degree of similarity between any two MRI volumes across all image analysis groups. With regard to ∆*FD/SSIM* pairs in within-subject comparisons, some cases were indicative of data clustering, and this was related to image registration. Figures 9 and 10 show the results for the exemplary group of T1 high-resolution images in the MASSIVE data set. In first volume registration, *k – means* clustering showed that the data was clearly separated into fractality-similarity clusters (panel A, fig. 9) across all analysis groups. Notably, this clustering was mainly driven by comparisons involving the first volume, i.e. the registration target (indexed by 0, see caption). Consequently, a number of across-cluster correlations were found in various analysis groups, suggesting a systematic negative association between differences in fractal dimension and structural similarity (panel B). However, this relationship was limited to clusters that were highly separated in both ∆*FD* and *SSIM* (cf. centroid location) and that were most clearly induced by comparisons involving the registration target. Indeed, when the same procedure was applied to high-resolution T1 images in the reregistered MASSIVE data, these associations disappeared (fig. 10). Here, ∆*FD/SSIM* clusters as found by *k − means* were generally less separated, mainly differed only by ∆*FD* in centroid location, and showed no systematic relationship between cluster assignment and which of the MRI volumes entered the comparison (panel A). Similarly, the previous associations between ∆*FD* and *SSIM* were strongly attenuated, and all but one vanished altogether (panel B). In fact, no general systematic relationship between fractal dimensions and structural similarity was observed in the reregistered MASSIVE data set. We then applied the same within-subject analysis to the MSC data. While we observed similar target-induced clustering and cluster-driven ∆*FD/SSIM* associations in first volume registration as well as the attenuation of these effects in the reregistered images (see appendix for an example), within-subject analyses in the MSC data set were restricted to only a few possible between-volumes comparisons due to the lower number of per-subject scans (cf. eq. 4). Nonetheless, the MSC data enabled us to compute extensive across-subject comparisons, as detailed in sec. 2.3.3. Figure 11 summarizes the results for the exemplary case of high-resolution T1 images, while similar results were found for T2WI. *k − means* clustering yielded two to three ∆*FD/SSIM* clusters for each image analysis group, with low between-cluster separation and centroid locations driven predominantly by differences in ∆*FD* or *SSIM* but not both (panel A). Furthermore, no systematic relationship between ∆*FD* and *SSIM* was observed for across-subjects comparisons (panel B).

**Figure 9:**
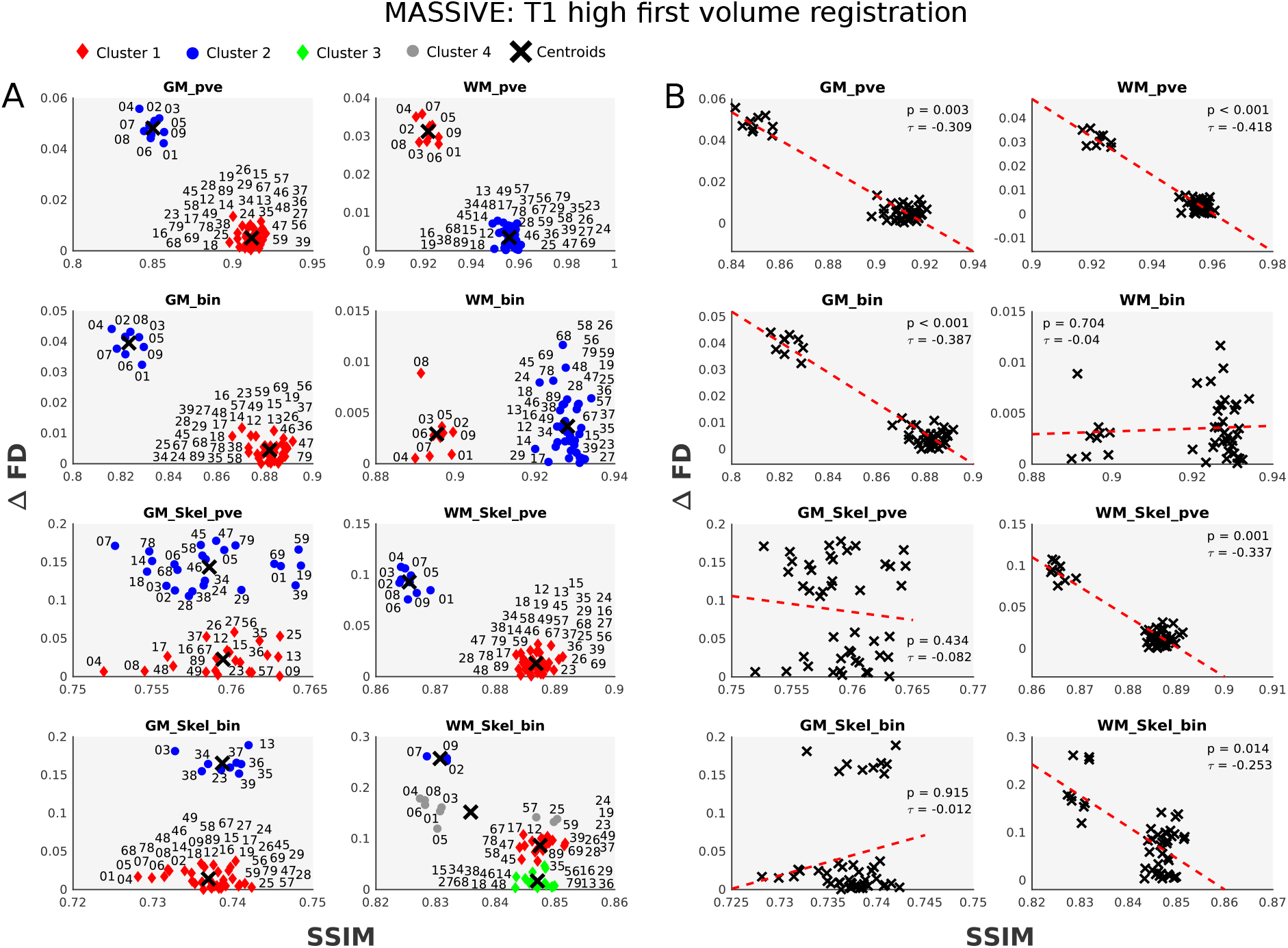
Fractal dimension differences and structural similarity in the MASSIVE high-resolution T1 images (first volume registration). Panel A displays the *k – means* clustering results within each analysis group. For all possible 45 comparisons, the Structural Similarity Index (*SSIM*) between two input volumes was computed and related to the difference in the corresponding fractal dimensions (∆*FD*). Numbers indicate which of the ten volumes were compared, with indices running from 0 to 9 to avoid triple digits. For first volume registration, ∆*FD/SSIM* pairs showed strong clustering, and there was a systematic effect of comparisons involving the first volume (the original registration target, indexed by 0) for most image analysis groups. In these groups, clusters were driven by differences in both ∆*FD* and *SSIM*, and this induced strong negative associations between differences in fractal dimension and structural similarity (panel B). This effect, however, was attenuated by reregistration (see fig. 10 and main text). ∆*FD*: absolute difference in fractal dimension between two compared volumes; GM: gray matter; bin: binary tissue segmentation; pve: partial volume estimates; Skel: skeleton model; *SSIM* Structural Similarity Index between two compared volumes; WM: white matter.

**Figure 10:**
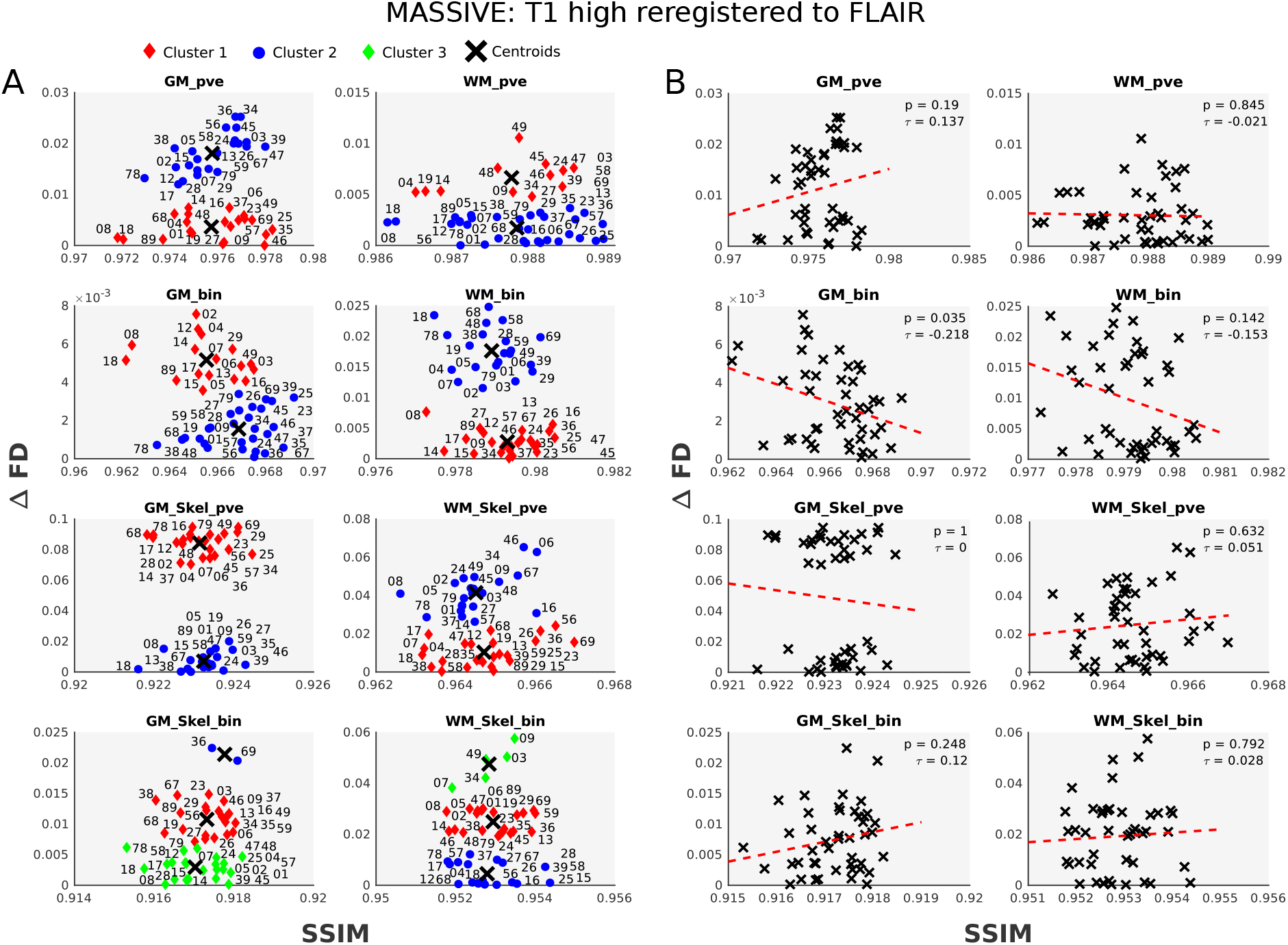
Fractal dimension differences and structural similarity in the MASSIVE high-resolution T1 images (reregistered to FLAIR). Similar to fig. 9, panel A represents the ∆*FD/SSIM* pairs and *k – means* clustering results for high-resolution T1 images after reregistration to FLAIR. Here, ∆*FD/SSIM* clusters as found by *k means* clustering were generally less separated, mainly differed only by ∆*FD* in centroid location, and showed no systematic relationship between cluster assignment and which of the input volumes entered the comparison (panel A). The previous associations between ∆*FD* and *SSIM* were strongly attenuated, and all but one vanished altogether (panel B). ∆*FD*: absolute difference in fractal dimension between two compared volumes; GM: gray matter; bin: binary tissue segmentation; pve: partial volume estimates; Skel: skeleton model; *SSIM* Structural Similarity Index between two compared volumes; WM: white matter.

**Figure 11:**
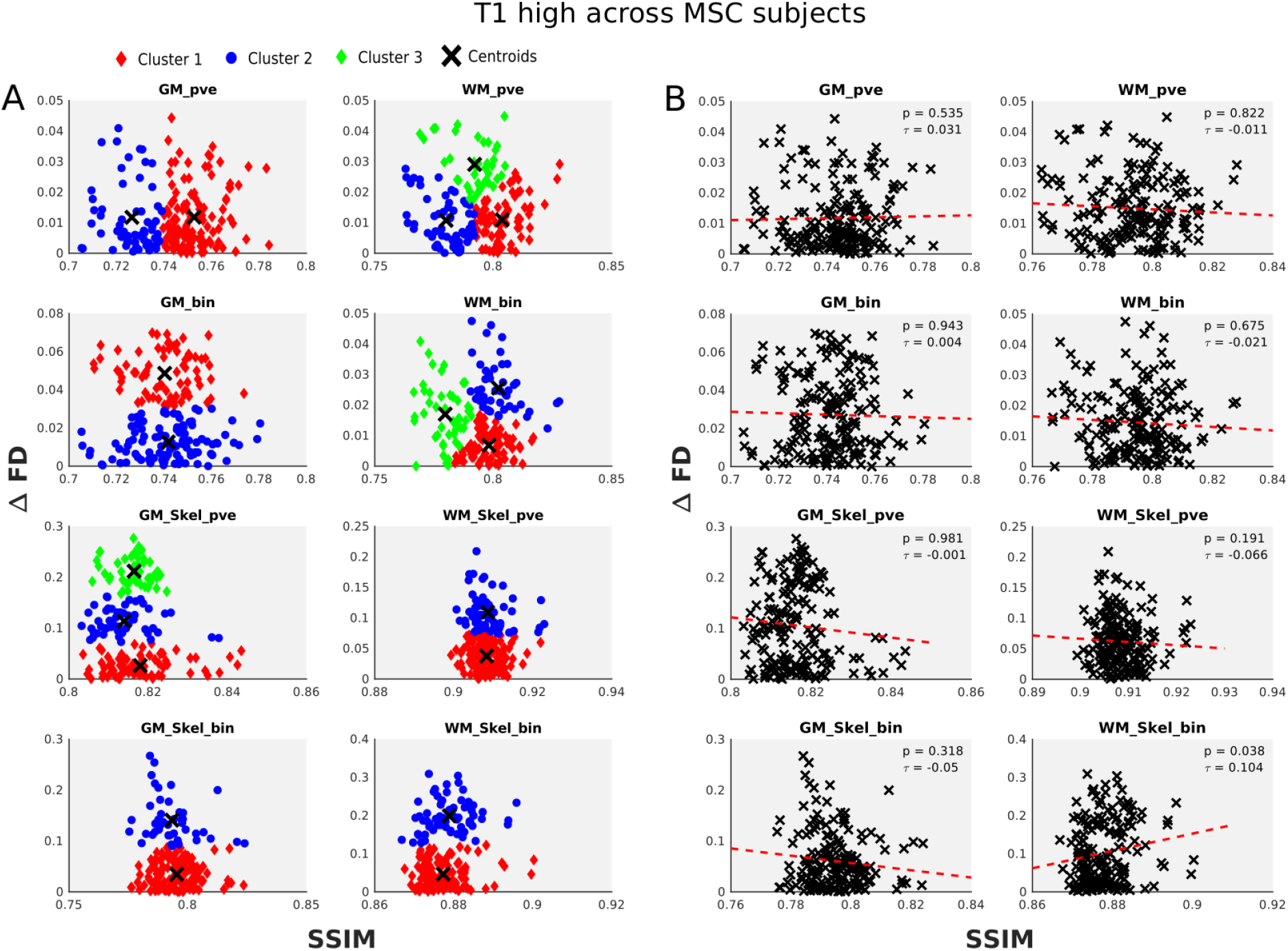
Fractal dimension differences and structural similarity in across-subjects comparisons in the MSC data set. Panel A visualizes the results of across-subject comparisons for high-resolution T1 images in MNI registration. Each subject had four scans, and all possible between-subject comparisons were computed for each of those scanning sessions across all image analysis groups (where we omit the comparison indices from above for visual coherence). Panel B relates the corresponding correlation results by image analysis groups. There was no systematic ∆*FD/SSIM* data clustering in across-subject comparisons, and no systematic association between the fractal dimension and structural similarity was found. GM: gray matter; bin: binary tissue segmentation; pve: partial volume estimates; Skel: skeleton model; *SSIM* Structural Similarity Index between two compared volumes; WM: white matter.

Further evidence against a systematic fractality-similarity association comes from between-registration comparisons of ∆*FD* and *SSIM* (see appendix). While *SSIM* values in the MASSIVE data set were significantly different between first volume registration and the reregistered data across all analysis groups (*p* < 0.001 for all comparisons, Bonferroni-Holm-adjusted), there was no significant difference in ∆*FD* values in the majority of the analysis groups (20/32 confirmed null hypotheses, cf. table I in appendix). This finding was corroborated and indeed more pronounced in the MSC data set, in which *SSIM* values for all analysis groups also showed a highly significant between-registration difference, while there were essentially no significant differences in ∆*FD* values between the two image registrations (30/32 confirmed null hypotheses, cf. table II in appendix).

#### 3.5 Fractal dimension by image characteristics

Finally, we compare the mean fractal dimension estimates by image weighting and resolution in a parameter-dependent fashion. Figure 12 reports the results for the MSC data set (but results for the MASSIVE data were highly similar, see fig. 3 in the appendix). As a general result, fractal dimension estimates in both image weighting were sampled in the expected range, compatible with previous reports, and T1WI and T2WI were affected by image registration, binarization, skeletonization, and spatial resolution in a highly similar manner. While the results from sec. 3.2 highlight that registration had a significant impact on the absolute fractal dimension values, the influence of sequence weighting, tissue type and image processing parameters within a given set of input images was essentially unaltered by reregistration. As such, binary tissue segmentation consistently caused a moderate reduction of FD values in the unskeletonized volumes across both registrations, while it led to a slight increase or no significant change in the skeleton models for both T1WI and T2WI, gray matter as well as white matter segmentations and regardless of spatial resolution. Furthermore, skeleton models invariably resulted in significantly decreased FD values across all analysis groups and in both image registrations. Another interesting pattern was observed with regard to tissue type: while gray matter and white matter fractal dimensions showed no significant differences for most comparisons in unskeletonized analysis groups, skeleton models generally yielded significantly higher gray matter fractal dimensions in T1WI as well as slightly but significantly higher white matter fractal dimensions for most comparisons in T2WI. Moreover, lower spatial resolution invariably resulted in significantly decreased fractal dimension values for all unskeletonized image groups in both the MASSIVE and the MSC data sets, regardless of image registration (see appendix, fig. 4). The same effect was observed in most image skeleton groups across both data sets, with a few exceptions in the MNI-registered MSC data. Furthermore, comparing the standard deviations in panels A and B of fig. 12, there was a marked reduction in between-subject variability by reregistration to MNI space for all unskeletonized analysis groups, while within- and between-subject variability were not equivalently reduced in skeleton models (cf. also by-subject averages in appendix, fig. 5).

**Figure 12:**
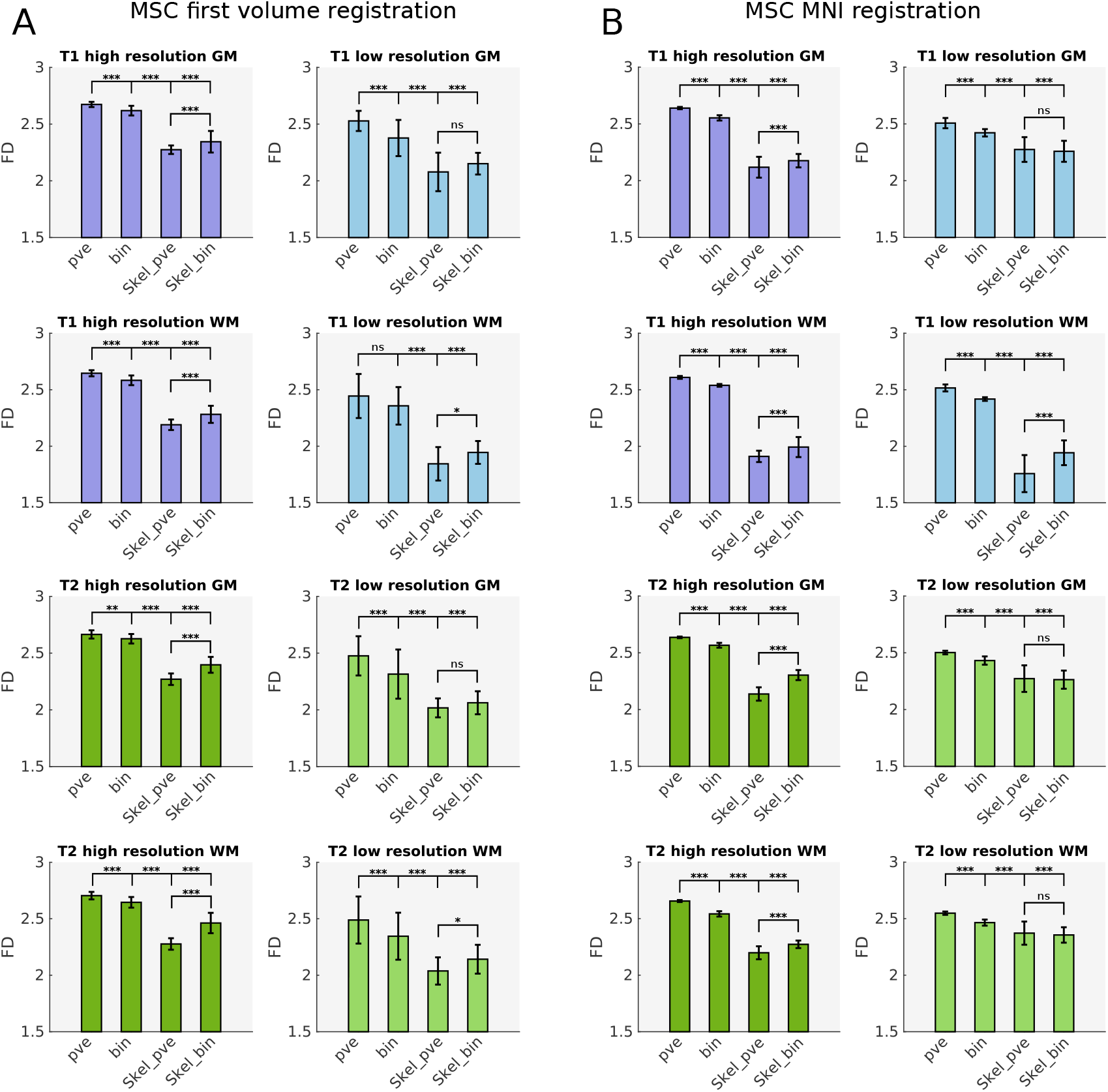
Parameter-dependent comparison of the fractal dimension estimates in the MSC data set. Panels A and B visualize the comparisons of the mean fractal dimension estimates over image analysis groups in first volume and MNI registration, respectively. Horizontal bars reflect pair-wise significance levels. Comparisons for binary-segmented images (second bar in each subpanel) invariably yielded the same significance levels as the partial volume estimates (first bar) so they were omitted here for visual coherence. Note that while image registration had a profound impact on the absolute fractal dimension estimates, the relative impact of sequence weighting, spatial resolution, segmentation procedure, tissue type, and skeletonization was essentially unaltered by registration. *ns*: not significant; *: *p <* 0.05; **: *p <* 0.01; ***: *p <* 0.001; bin: binary tissue segmentation; GM: gray matter; pve: partial volume estimates; Skel: skeleton model; WM: white matter.

## 4 Discussion

The current study presents a systematic and in-depth evaluation of the fractal dimension as a marker of structural brain complexity in human brain MRI. To this end, we first consider some computational aspects regarding fractal dimension estimation based on 3D box-counting, and we report a detailed empirical analysis of two recently published open-access neuroimaging data sets.

As detailed above, the fractal dimension estimates obtained from box-counting numerically depend on the spatial scale interval over which the linear regression of the log-transformed data is computed, highlighting the question which scale interval will most adequately capture the underlying fractal dimension in the estimation process. We here applied an algorithmic scale optimization procedure to address this issue. The outlined procedure led to a significant improvement of estimation accuracy over randomly chosen non-optimal scales, as suggested by simulation studies of random Cantor sets whose expected fractal dimension values were known. Interestingly, these results also indicated that performance against optimization was not uniform across all non-optimal spatial scales, i.e. that under- and overestimation of the true fractal dimension varied from moderate to severe, depending on how non-optimal a randomly chosen control interval was. This finding illustrates that choosing inadequate spatial intervals for fractality estimation may entail the danger of pronounced and systematic inaccuracies, potentially obscuring utility in empirical estimation. In this regard, we suggest that the applied procedure provides improvement over using fixed spatial scales, as is commonly done. In similar spirit, group-wise scale selection based on correlation maximization has been applied by Esteban et al. 2010. Nonetheless, generalization of optimal scales across distinct estimations may be limited by differences in populations, subjects, acquisition sessions, scanning equipment, or estimation software, and thus a more data-driven approach offers increasingly individualized optimization. In the current study, we apply scale optimization to individual fractality estimations in a completely automatic fashion.

This procedure also enabled us to analyze which spatial scales were in fact selected as optimal from the empirical data and relate optimization results to subjects and image characteristics. Indeed, the results from sec. 3.3 show that optimal *k*-ranges were selective in terms of interval length, numerical *k*-values (i.e. box edge sizes) and image analysis groups. Here, an interesting pattern emerged in function of skeletonization, with remarkably similar optimization outcomes in the MASSIVE and the MSC data sets and high consistency across repeated measurements and subjects. As such, it will be interesting to see in future studies whether this represents a general property of the box-counting estimation in brain MRI or if and to what extent it is specific to other factors, such as the preprocessing software, the examined set of spatial scales, or the applied optimization decision criterion. With regard to the latter, scale selection here was based on the adjusted coefficient of determination, a commonly used measure of goodness of fit. While this perhaps represents the most natural approach to the box-counting regression, other well-studied model selection criteria exist (e.g. the Bayesian Information Criterion), and future studies may examine if applying a different model selection criterion yields further improvement of estimation accuracy. In this context, it is also interesting to note that the disadvantage of using all spatial scales in the box-counting regression has been pointed out from an analytical perspective (cf. e.g. Gneiting et al. 2012), and indeed avoidance of greater-length *k*-ranges was observed in our empirical optimization outcomes. Finally, further study is also warranted to examine if similar improvements can be achieved in other methods of fractality estimation, such as dilation-based algorithms, which are thought to possess several advantages over classical box-counting (e.g. Madan and Kensinger 2016, 2017).

With regard to our empirical results, deviation analyses suggested a high overall test-retest stability of the fractal dimension estimates (approximately 95 %) across both the MASSIVE and the MSC data sets. This is in accordance with a recent reliability study of brain morphology estimates in two open-access data sets by Madan and Kensinger (Madan and Kensinger, 2017) who found that regional fractal dimensionality as computed by both dilation and box-counting methods was generally very high and comparable to the reliability of gyrification indices, while it was in fact superior to volumetric measures such as cortical thickness. Similarly, Goñi and colleagues analyzed the fractal properties of the pial surface, the gray matter / white matter boundary and the cortical ribbon and white matter volumes in MRI data from different imaging centers and found a high within-subject reproducibility with region-specific patterns of individual variability (Goñi et al., 2013). While there is thus converging evidence for the robustness of fractal analysis in neuroimaging, these studies used parcellation- and surface-based methods, and T2WI were not analyzed. In this regard, the present study provides additional information as our evaluation was stratified into 32 distinct analysis groups based on sequence weighting, spatial resolution, segmentation procedure, tissue type, and image complexity, highlighting that the different image variables entail a differential susceptibility to repeated-sampling deviations, observed here especially for high-resolution images and skeleton models.

In this context, one important finding of the current study concerns the complex and profound influence of image registration on the fractal dimension estimates. In both data sets, image registration had a significant impact on the absolute fractal dimension estimates, without obvious patterns across the various analysis groups. Furthermore, we found that unbalanced registration targets can induce test-retest deviations in the fractal dimension estimates that are reduced with reregistration, and this was consistently observed in both data sets and across subjects in the MSC data. These test-retest deviations were also found to render the assumption of composite normality to be invalid in repeated within-subject sampling. While a high proportion of analysis groups in balanced registration adhered to composite normality for repeated within-subject measurements, this did not transfer to the across-subject sample distributions. Instead, here the assumption of normality was refuted in a large majority of image analysis groups, and differences between analysis groups appeared to be driven by a variable across-subject sample regularization in balanced registration. This finding (together with the test-inherent limitation that accepting the null hypothesis does not prove composite normality but rather indicates it should not be refuted) suggests that it may not be advisable to assume the fractal dimension estimates over various subjects to be sampled from an underlying normal distribution. Measuring multiple subjects with only one or a few respective samples is a very common empirical scenario, however. As such, it appears that distributional assumptions in comparisons across populations (e.g. patients vs. controls) may need to be relaxed, for instance by opting for non-parametric methods, or ought to be informed by explicit assessment.

Furthermore, image registration also had an interesting effect on between-group ties within the data sets: while unbalanced registration induced strong associations among various analysis groups, reregistration caused a pronounced overall decorrelation (indeed, the presence of strong across-groups associations also seems biologically implausible, e.g. there is no principled reason to believe that structural complexity of white matter will generally follow that of gray matter). In summary, our results point to an important methodological question: given the profound impact of image registration of the fractality estimates, which registration scheme should be applied for fractal analysis of structural brain MRI? While our results clearly argue for balanced registration methods, it is at this point unclear if subject-derived templates (that were found to increase between-scan structural similarity, see below) carry any advantages over subject-independent templates. In any case, as the former may not always be feasible (e.g. in single-acquisition scenarios), registration to commonly used subject-independent targets such as the MNI template may currently be a reasonable solution, perhaps also in the interest of between-study comparisons.

The latter point also concerns procedural standardization and technical variance. Both the MASSIVE and the MSC data set provide highly standardized images, while this may not always be the case in empirical reality. Motion artifacts, for instance, can be expected to obscure the utility of fractal analysis. Similarly, just as reference values for blood tests may differ depending on the laboratory where they are measured, fractal analysis may be influenced by the type of scanning equipment, sequences, preprocessing software or estimation method, as has been shown for other morphometric analyses, e.g. Wonderlick et al. 2009; Madan and Kensinger 2017; Duché et al. 2017. In this context, however, it is noteworthy that the MASSIVE and the MSC data were acquired on scanning systems from two different manufacturers. While this provides some evidence that the results presented herein (which were very similar across the two data sets) were fairly independent of the scanning equipment, it may also constitute one reason why the absolute numerical dimension estimates were not generally transferable from one data set to another.

One finding with high consistency across the two data sets regards the impact of binary segmentation on the fractal dimension estimates, which caused a moderate FD reduction in unskeletonized images but no change or slight increases in skeleton models. In this context, it is noteworthy that image skeletons invariably yielded decreased fractality estimates as compared to their unskeletonized counterparts across both data sets. Since the skeleton models can be thought of as a minimum complexity version of the input volume, it seems rather plausible that the fractal dimension as a marker of tissue complexity was consistently reduced by skeletonization. Furthermore, it is interesting to note that the lower voxel resolution invariably resulted in lower FD values in the unskeletonized images across both data sets. For skeletonized images, a similar pattern was observed, with a few exceptions in the MNI-registered MSC data set. Intuitively, a measure of structural brain complexity may be decreased in coarser spatial resolution because structural information is blunted by partial volume effects.

Furthermore, the present study systematically evaluates the methodological characteristics of fractality estimation in structural T2WI. While T2-derived sequences have been used for fractal analysis in the realm of functional MRI (albeit predominantly with respect to time series analysis, e.g. Bullmore et al. 2001; Thurner et al. 2003; Foss et al. 2006; Lai et al. 2010; Eke et al. 2012), T1WI have been the mainstay of structural neuroimaging studies employing fractal analysis. Nonetheless, there has been some prior indication that T2-based fractal analysis is both feasible and useful, especially in clinical assessment. For instance, Iftekharuddin and colleagues successfully incorporated T2WI in fractality-based multimodal feature extraction for tumor segmentation (Iftekharuddin et al., 2009), and Takahashi et al. used multi-fractal analysis of deep white matter in T2WI to detect microstructural changes in early atherosclerotic alterations (Takahashi et al., 2006). Furthermore, Di Ieva and colleagues characterized nidus angioarchitecture of brain arteriovenous malformations with fractal analysis of T2WI (Di Ieva et al., 2014b). In the present study, we found T2WI to yield remarkably robust results, both in comparison to T1WI and in terms of stability over repeated measurements. Furthermore, T1WI and T2WI were affected by binarization, skeletonization, and spatial resolution in a similar manner, which may encourage further research given the importance of T2WI in clinical neuroradiological practice.

Finally, perhaps one of the most interesting findings of this study concerns the relationship between structural complexity and structural similarity. These analyses were motivated both by registration-induced changes and by the general question of whether differences in fractal dimension essentially just reflect differences in structure per se. To our knowledge, the present study is the first to investigate the relationship between the fractal dimension and the structural similarity index (*SSIM*) in MRI.

Structural similarity as captured by the *SSIM* was generally very high across the two data sets. In relating structural similarity to the corresponding difference in fractal dimensions (∆*FD*), we applied a *k − means* clustering analysis, which provided a useful way to objectively assess data clusters, especially since *k* was chosen automatically and the same optimization settings were used for both image registrations and across both data sets. Due to the method’s unsupervised character, it can be difficult to interpret qualitative differences in the clusters’ features. However, based on the procedure in sec. 2.3.3, each ∆*FD/SSIM* pair represented a particular comparison of two MRI volumes, thus enabling us to check for systematic effects of between-volume comparisons as cluster features, and comparing the cluster centroids was useful in describing whether clustering was mostly driven by differences in ∆*FD*, *SSIM*, or both. We furthermore conducted the analyses in two distinct ways: we first examined fractality-similarity relationships over repeated acquisitions within subjects, and then extended the analyses to comparisons across subjects.

In line with the results of sec. 3.2, we found considerable within-subject clustering in various analysis groups for the MASSIVE data set in first volume registration, with a systematic effect of comparisons involving the registration target that yielded pronounced between-cluster separation in both ∆*FD* and *SSIM* and induced a number of strong fractality-similarity correlations. However, we interpret these to be spurious correlations induced by unbalanced registration because 1) they were mostly limited to analysis groups with strong target-induced clustering, 2) the direction of the association was not consistent across analysis groups, 3) structural similarity across all analysis groups was significantly different in the reregistered data set (as expected) while there was little difference in ∆*FD*, and 4) systematic ∆*FD/SSIM* clustering and fractality-similarity associations essentially disappeared with reregistration. While we observed a similar tendency in the MSC data set towards target-induced clustering entailing across-cluster associations in first volume registration and the attenuation thereof in MNI registration, within-subject comparisons were numerically limited by the lower number of per-subject scans as compared to the MASSIVE data. However, the MSC data allowed for extensive across-subject comparisons, which showed no systematic ∆*FD/SSIM* clustering and no association between fractal dimension differences and structural similarity. Furthermore, similar to the MASSIVE data, structural similarity across all image groups was significantly different between first volume and MNI registration, while there was essentially no difference in ∆*FD*. In this context, a closer examination of the numerical *SSIM* values in the MASSIVE and the MSC data reveals a subtle but interesting corollary of our analyses: while reregistration in the MASSIVE data set invariably caused a marked increase in the *SSIM* values to above 0.9 in all analysis groups, reregistration in the MSC data caused a decrease in *SSIM* in all but two analysis groups to values around 0.7 - 0.8 (cf. tables I and II in the appendix). Bearing in mind that the MASSIVE data were reregistered to the mean FLAIR image (derived from the same subject) while the MSC data were reregistered to the MNI template (i.e. not derived from the same subjects), these findings suggest that subject-specific common image registration increased between-scan structural similarity while subject-independent common registration decreased between-scan similarity. Notably, however, differences in fractal dimension did not simply track differences in structural similarity in either case, i.e. regardless of whether scans were more or less similar to each other, and this applied to both within- and across-subject analyses. In summary, the present results suggest that there is no general relationship between structural complexity as measured by the fractal dimension and structural similarity as captured by the structural similarity index and that, rather, they may represent two distinct aspects of structural brain MRI.

### 4.1 Future directions

In the current study, we obtain several fractal dimension values for every input volume due to the stratification of processing parameters (tissue type, segmentation procedure, image complexity). Thus, instead of just mapping one fractal dimension to one image, we compute a fractal "profile" of eight fractal dimension estimates per input image. Since the different analysis groups seem to entail differential susceptibility to deviations, such a fractal profile could perhaps be useful to optimize diagnostic sensitivity-specificity trade-offs. Furthermore, we used a monofractal analysis approach, and it may be useful to expand this to multifractal analysis. Moreover, we here compute fractal dimension estimates on global tissue segmentations. Given the increasingly sophisticated brain parcellation methods, however, region-and substructure-specific fractal analysis is also being developed and is likely to yield interesting additional information, especially in the clinical context (see e.g. Goñi et al. 2013; Glasser et al. 2016; Madan and Kensinger 2017; Ruiz de Miras et al. 2017; Madan 2018).

## 5 Acknowledgments

The work of S.K. and C.F. is funded by the German Federal Ministry for Education and Research (BMBF grant 13GW0206D). The research of A.L. is supported by VIDI Grant 639.072.411 from the Netherlands Organization for Scientific Research (NWO). P.V. is employed by Genentech Inc for work unrelated to this project. P.V. has stocks and is serving in the advisory board of Health Engineering SL (who has licensed a platform for measuring the fractal dimension from brain images from the University of Jaén and IDIBAPS), QMenta SL and Bionure SL. The work of F.J.E. is supported by Junta de Andalucía (BIO-302) and MEIC (Systems Medicine Excellence Network SAF2015-70270-REDT). Finally, we are grateful to Jakob Ludewig and Leonhard Waschke for inspiring discussions regarding the current work.

## Appendix

The following provides some supporting material underlining aspects from the main text. While we here limit ourselves to a selection of some relevant illustrations, the interested reader is encouraged to review our implementation for further exploration of the method and the data sets (see http://osf.io/3mtqx). The implementation package also includes a demo function that reiterates the presented analyses, creating figures and tables from the main text and the appendix.

Below, fig. 1 displays the silhouette optimization results for the across-subject ∆*FD/SSIM k*- means analysis in the Midnight Scan Club (MSC) data set from the main text. Clustering analysis was implemented with the Statistics and Machine Learning Toolbox for Matlab. We compared various ranges of *k* over which to carry out silhouette optimization, with the range [1, 4] showing highest consistency across image groups in terms of finding the same clusters compared to other ranges while avoiding overclustering. For silhouette optimization, each cluster contributed to the overall silhouette values in proportion to its size. The squared euclidean distance was used as the metric for point-to-centroid distance minimization. Initial centroid positions were chosen randomly from the data set, i.e. the *k − means* ++ heuristic was used to optimize time to convergence. We applied the clustering algorithm over ten replicate iterations in every case to avoid convergence on non-global distance minima. Importantly, the same settings for silhouette optimization and clustering were applied to both the MASSIVE and the MSC data sets and in both image registrations.

Furthermore, fig. 2 displays an exemplary evaluation of within-subject ∆*FD/SSIM k*-means analysis for high-resolution T1-weighted images of the first subject in the MSC data set, equivalent to figures 9 and 10 in the MASSIVE data from the main text. While there was a similar tendency towards target-induced ∆*FD/SSIM* clustering entailing across-cluster associations in first volume registration as well as the attenuation of these effects by reregistration, here the number of possible within-subject comparisons was limited to 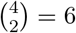 as opposed to 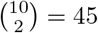 in the MASSIVE data set.

Moreover, tables I and II relate the between-registration comparisons of the mean ∆*FD* and *SSIM* values for the MASSIVE and the MSC data sets, respectively. While image registration led to a significant change in *SSIM* values in both data sets, there was no equivalent change in ∆*FD* values. Interestingly, registering images to a subject-derived target invariably augmented structural similarity (MASSIVE, table I), while a common but subject-independent registration target generally decreased structural similarity (MSC, table II).

In addition, figure 3 displays the parameter-dependent comparisons of the mean fractal dimension estimates in the MASSIVE data set, in analogy to the results presented for the MSC data set in figure 12 of the main text. There was a remarkable similarity between the two data sets regarding the impact of sequence weighting, binarization, tissue type, and skeletonization. Image registration, while significantly affecting the absolute fractal dimension values (cf. table I in the main text), had little effect on the relative influence of these analysis parameters (e.g. binarization affected FD estimates in a similar way in both registrations, etc.), and this was equivalently observed in both the MASSIVE and the MSC data. Similarly, the effect of spatial resolution was quite consistent across the two data sets and is summarized in figure 4. Lower spatial resolution invariably resulted in decreased fractal dimension estimates for unskeletonized images across both data sets and registrations. A similar pattern was observed for skeleton models, with a few exceptions in the MNI-registered MSC data set.

Furthermore, figure 5 reports the by-subject average fractal dimension estimates across both image registrations in the MSC data set. Between-subject variability was higher in first volume as compared to MNI registration for unskeletonized image groups, while both between-subject and within-subject variability were camparatively high in skeleton models for either image registration.

**Figure 1:**
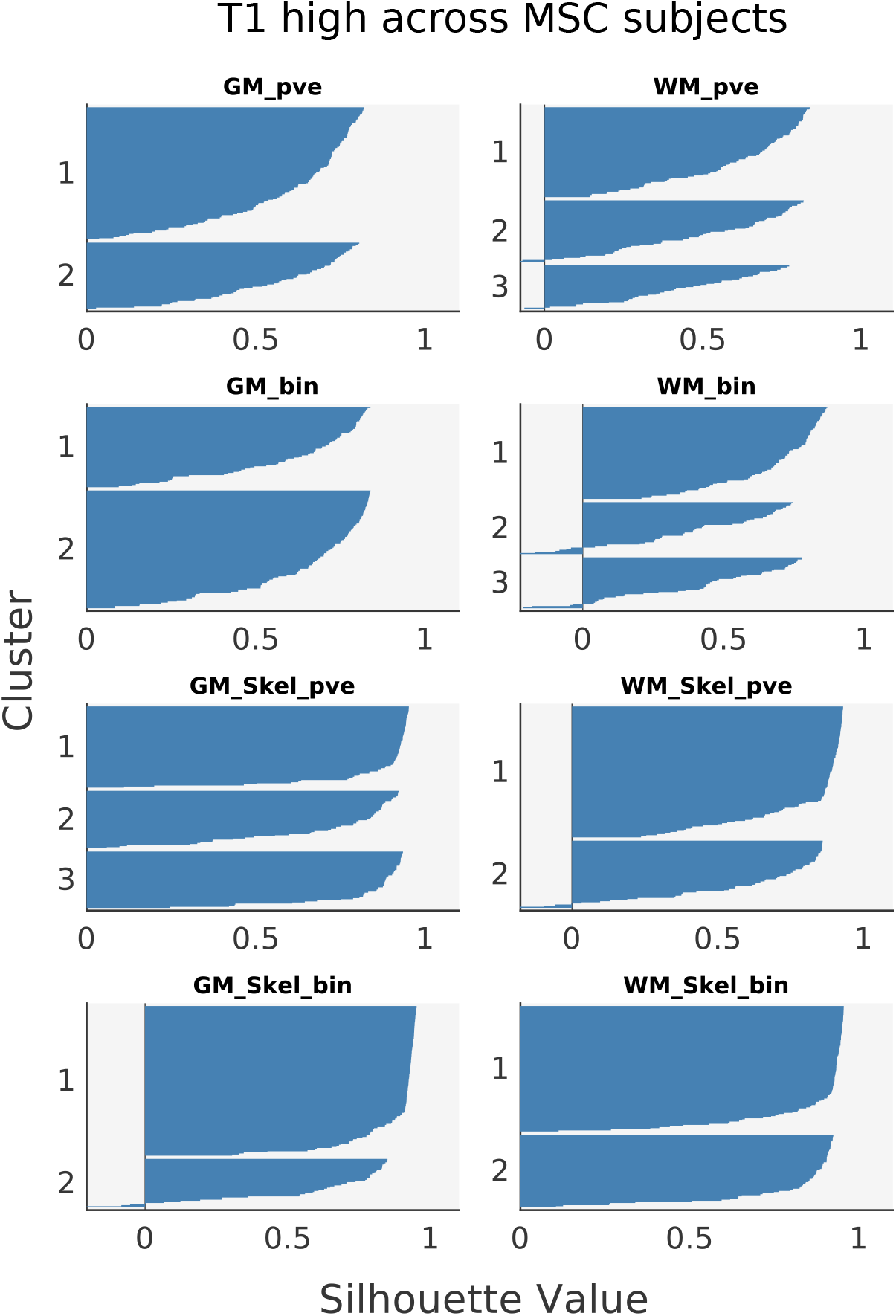
Silhouette plots for *k – means* clustering analysis of ∆*FD/SSIM* pairs in across-subject comparisons (MSC). The figure relates the silhouette values belonging to the exemplary across-subject comparisons in the T1 high-resolution images from the MSC data set, as presented in the main text. Estimation outcome predominantly features positive observations, with most values ranging between 0.7 and 1, indicative of good clustering quality. The optimal number of clusters was determined automatically by silhouette optimization, given here by *k* = 2 in five analysis groups and *k* = 3 in the remaining three cases. Cluster sizes were reasonably balanced in most instances, and there was no indication of overclustering. GM: gray matter; bin: binary tissue segmentation; pve: partial volume estimates; Skel: skeleton model; WM: white matter.

**Figure 2:**
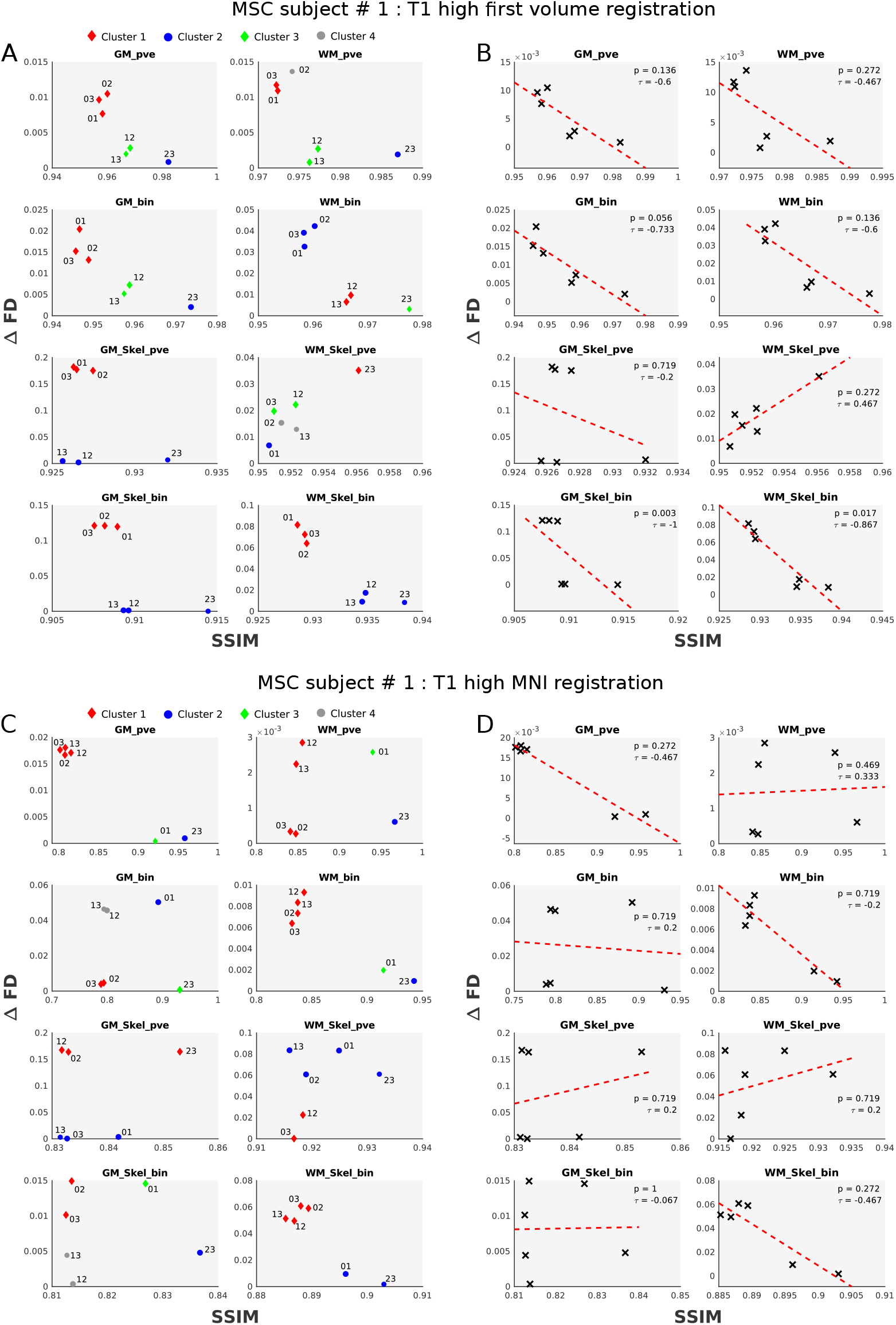
Exemplary within-subject analysis of fractal dimension and structural similarity in the MSC data set. Panel A relates the *k – means* clustering of ∆*FD/SSIM* pairs for all possible within-subject comparisons in the high-resolution T1 images of the first subject in the MSC data set. There was an indication of target-induced clustering (panel A) entailing across-cluster associations (panel B) in first volume registration as well as the attenuation thereof in MNI registration (panels C and D), similar to the findings in the MASSIVE data. However, as there were only four repeated acquisitions, the number of possible within-subject comparisons was limited to 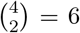, as opposed to 45 per analysis group in the MASSIVE data. GM: gray matter; bin: binary tissue segmentation; pve: partial volume estimates; SSIM: structural similarity index; WM: white matter.

**Table I:** Between-registration comparison of fractal dimension differences and structural similarity in the MASSIVE data set. The table represents the mean ∆*FD* and *SSIM* values in the first volume and FLAIR-registered data set compared by the Wilcoxon rank sum test. All p-values are Bonferroni-Holm-corrected for multiple comparisons (*p_corr_*). GM: gray matter; bin: binary tissue segmentation; pve: partial volume estimates; Skel: skeleton model; SSIM: structural similarity index; WM: white matter.

| Analysis group | $\Delta FD_{first}$ | $\Delta FD_{FLAIR}$ | $p_{corr}$ | $SSIM_{first}$ | $SSIM_{FLAIR}$ | $p_{corr}$ |
| --- | --- | --- | --- | --- | --- | --- |
| **T1 high** |  |  |  |  |  |  |
| GM_pve | $0.0137 \pm 0.0178$ | $0.0113 \pm 0.0079$ | *ns* | $0.8994 \pm 0.0256$ | $0.9757 \pm 0.0015$ | $1.0e-14$ |
| GM_bin | $0.0115 \pm 0.0145$ | $0.0029 \pm 0.0020$ | 0.0109 | $0.8702 \pm 0.0244$ | $0.9664 \pm 0.0016$ | $7.0e-15$ |
| GM_Skel_pve | $0.0871 \pm 0.0642$ | $0.0482 \pm 0.0395$ | 0.0173 | $0.7590 \pm 0.0031$ | $0.9232 \pm 0.0006$ | $1.0e-14$ |
| GM_Skel_bin | $0.0442 \pm 0.0622$ | $0.0074 \pm 0.0054$ | 0.0023 | $0.7373 \pm 0.0032$ | $0.9172 \pm 0.0007$ | $8.9e-15$ |
| WM_pve | $0.0090 \pm 0.0115$ | $0.0030 \pm 0.0025$ | *ns* | $0.9490 \pm 0.0140$ | $0.9878 \pm 0.0007$ | $8.6e-15$ |
| WM_bin | $0.0035 \pm 0.0027$ | $0.0097 \pm 0.0080$ | 0.0227 | $0.9216 \pm 0.0137$ | $0.9791 \pm 0.0008$ | $8.2e-15$ |
| WM_Skel_pve | $0.0289 \pm 0.0335$ | $0.0249 \pm 0.0179$ | *ns* | $0.8826 \pm 0.0088$ | $0.9646 \pm 0.0009$ | $9.8e-15$ |
| WM_Skel_bin | $0.0862 \pm 0.0696$ | $0.0193 \pm 0.0147$ | $1.4e-05$ | $0.8439 \pm 0.0074$ | $0.9529 \pm 0.0007$ | $7.6e-15$ |
| **T1 low** |  |  |  |  |  |  |
| GM_pve | $0.0093 \pm 0.0065$ | $0.0075 \pm 0.0056$ | *ns* | $0.8916 \pm 0.0205$ | $0.9704 \pm 0.0034$ | $6.7e-15$ |
| GM_bin | $0.0170 \pm 0.0133$ | $0.0046 \pm 0.0031$ | 0.0038 | $0.8590 \pm 0.0205$ | $0.9585 \pm 0.0039$ | $7.9e-15$ |
| GM_Skel_pve | $0.0086 \pm 0.0058$ | $0.0228 \pm 0.0303$ | *ns* | $0.6975 \pm 0.0121$ | $0.9032 \pm 0.0027$ | $6.3e-15$ |
| GM_Skel_bin | $0.0107 \pm 0.0123$ | $0.0090 \pm 0.0068$ | *ns* | $0.6994 \pm 0.0086$ | $0.9045 \pm 0.0030$ | $6.0e-15$ |
| WM_pve | $0.0032 \pm 0.0024$ | $0.0039 \pm 0.0025$ | *ns* | $0.9524 \pm 0.0102$ | $0.9854 \pm 0.0019$ | $7.3e-15$ |
| WM_bin | $0.0129 \pm 0.0109$ | $0.0054 \pm 0.0035$ | *ns* | $0.9239 \pm 0.0098$ | $0.9754 \pm 0.0025$ | $9.5e-15$ |
| WM_Skel_pve | $0.0391 \pm 0.0308$ | $0.0570 \pm 0.0356$ | *ns* | $0.7946 \pm 0.0124$ | $0.9366 \pm 0.0027$ | $5.7e-15$ |
| WM_Skel_bin | $0.0264 \pm 0.0165$ | $0.1072 \pm 0.0875$ | 0.0022 | $0.7643 \pm 0.0090$ | $0.9236 \pm 0.0035$ | $2.5e-15$ |
| **T2 high** |  |  |  |  |  |  |
| GM_pve | $0.0020 \pm 0.0012$ | $0.0022 \pm 0.0014$ | *ns* | $0.9210 \pm 0.0198$ | $0.9779 \pm 0.0058$ | $5.4e-15$ |
| GM_bin | $0.0052 \pm 0.0034$ | $0.0051 \pm 0.0033$ | *ns* | $0.8948 \pm 0.0215$ | $0.9707 \pm 0.0059$ | $5.1e-15$ |
| GM_Skel_pve | $0.0143 \pm 0.0142$ | $0.0176 \pm 0.0116$ | *ns* | $0.7817 \pm 0.0118$ | $0.9348 \pm 0.0034$ | $4.8e-15$ |
| GM_Skel_bin | $0.0268 \pm 0.0179$ | $0.0333 \pm 0.0291$ | *ns* | $0.7475 \pm 0.0134$ | $0.9252 \pm 0.0043$ | $2.2e-15$ |
| WM_pve | $0.0028 \pm 0.0019$ | $0.0042 \pm 0.0028$ | *ns* | $0.9037 \pm 0.0202$ | $0.9735 \pm 0.0055$ | $4.4e-15$ |
| WM_bin | $0.0066 \pm 0.0093$ | $0.0022 \pm 0.0016$ | *ns* | $0.8707 \pm 0.0239$ | $0.9648 \pm 0.0064$ | $1.9e-15$ |
| WM_Skel_pve | $0.0379 \pm 0.0248$ | $0.0190 \pm 0.0121$ | 0.0023 | $0.7473 \pm 0.0134$ | $0.9242 \pm 0.0041$ | $1.6e-15$ |
| WM_Skel_bin | $0.0130 \pm 0.0132$ | $0.0064 \pm 0.0048$ | *ns* | $0.7183 \pm 0.0166$ | $0.9169 \pm 0.0048$ | $9.2e-15$ |
| **T2 low** |  |  |  |  |  |  |
| GM_pve | $0.0016 \pm 0.0011$ | $0.0104 \pm 0.0136$ | $2.6e-06$ | $0.9301 \pm 0.0226$ | $0.9784 \pm 0.0072$ | $1.3e-15$ |
| GM_bin | $0.0012 \pm 0.0008$ | $0.0052 \pm 0.0038$ | $2.0e-07$ | $0.9046 \pm 0.0236$ | $0.9711 \pm 0.0072$ | $4.1e-15$ |
| GM_Skel_pve | $0.0177 \pm 0.0188$ | $0.0525 \pm 0.0776$ | *ns* | $0.6991 \pm 0.0291$ | $0.9090 \pm 0.0087$ | $9.5e-16$ |
| GM_Skel_bin | $0.0041 \pm 0.0030$ | $0.0089 \pm 0.0060$ | 0.0037 | $0.6894 \pm 0.0278$ | $0.9055 \pm 0.0077$ | $3.8e-15$ |
| WM_pve | $0.0056 \pm 0.0040$ | $0.0059 \pm 0.0037$ | *ns* | $0.9240 \pm 0.0213$ | $0.9769 \pm 0.0060$ | $3.5e-15$ |
| WM_bin | $0.0012 \pm 0.0009$ | $0.0061 \pm 0.0041$ | $1.3e-08$ | $0.8800 \pm 0.0270$ | $0.9638 \pm 0.0082$ | $3.2e-15$ |
| WM_Skel_pve | $0.0158 \pm 0.0134$ | $0.0493 \pm 0.0631$ | *ns* | $0.6522 \pm 0.0261$ | $0.8921 \pm 0.0076$ | $2.9e-15$ |
| WM_Skel_bin | $0.0037 \pm 0.0031$ | $0.0024 \pm 0.0016$ | *ns* | $0.6650 \pm 0.0315$ | $0.8974 \pm 0.0093$ | $6.3e-16$ |

**Table II:** Between-registration comparison of fractal dimension differences and structural similarity in the MSC data set. The table represents the mean ∆*FD* and *SSIM* values in the first volume and MNI-registered data set compared by the Wilcoxon rank sum test. All p-values are Bonferroni-Holm-corrected for multiple comparisons (*p_corr_*). GM: gray matter; bin: binary tissue segmentation; pve: partial volume estimates; Skel: skeleton model; SSIM: structural similarity index; WM: white matter.

| Analysis group | $\Delta FD_{first}$ | $\Delta FD_{MNI}$ | $p_{corr}$ | $SSIM_{first}$ | $SSIM_{MNI}$ | $p_{corr}$ |
| --- | --- | --- | --- | --- | --- | --- |
| **T1 high** |  |  |  |  |  |  |
| GM_pve | $0.0057 \pm 0.0071$ | $0.0077 \pm 0.0090$ | *ns* | $0.9475 \pm 0.0165$ | $0.8516 \pm 0.0700$ | $2.4e - 14$ |
| GM_bin | $0.0139 \pm 0.0121$ | $0.0156 \pm 0.0166$ | *ns* | $0.9240 \pm 0.0226$ | $0.8332 \pm 0.0609$ | $9.8e - 15$ |
| GM_Skel_pve | $0.0231 \pm 0.0368$ | $0.0616 \pm 0.0617$ | 0.0096 | $0.8778 \pm 0.0253$ | $0.8389 \pm 0.0114$ | $8.4e - 19$ |
| GM_Skel_bin | $0.0868 \pm 0.0920$ | $0.0468 \pm 0.0458$ | *ns* | $0.8497 \pm 0.0327$ | $0.8226 \pm 0.0145$ | $1.9e - 08$ |
| WM_pve | $0.0061 \pm 0.0049$ | $0.0057 \pm 0.0072$ | *ns* | $0.9612 \pm 0.0119$ | $0.8839 \pm 0.0560$ | $9.3e - 15$ |
| WM_bin | $0.0128 \pm 0.0128$ | $0.0082 \pm 0.0073$ | *ns* | $0.9350 \pm 0.0194$ | $0.8691 \pm 0.0468$ | $7.0e - 14$ |
| WM_Skel_pve | $0.0494 \pm 0.0482$ | $0.0368 \pm 0.0363$ | *ns* | $0.9111 \pm 0.0250$ | $0.9232 \pm 0.0071$ | $3.6e - 05$ |
| WM_Skel_bin | $0.0896 \pm 0.0750$ | $0.0547 \pm 0.0553$ | *ns* | $0.8823 \pm 0.0296$ | $0.8952 \pm 0.0086$ | $3.6e - 05$ |
| **T1 low** |  |  |  |  |  |  |
| GM_pve | $0.0211 \pm 0.0719$ | $0.0196 \pm 0.0489$ | *ns* | $0.9605 \pm 0.0178$ | $0.8388 \pm 0.1048$ | $1.7e - 17$ |
| GM_bin | $0.0158 \pm 0.0399$ | $0.0196 \pm 0.0317$ | *ns* | $0.9448 \pm 0.0215$ | $0.8018 \pm 0.0984$ | $7.2e - 19$ |
| GM_Skel_pve | $0.0397 \pm 0.0524$ | $0.0916 \pm 0.0974$ | *ns* | $0.8367 \pm 0.0507$ | $0.7311 \pm 0.0333$ | $7.1e - 20$ |
| GM_Skel_bin | $0.0245 \pm 0.0259$ | $0.0524 \pm 0.0677$ | *ns* | $0.8216 \pm 0.0447$ | $0.7163 \pm 0.0319$ | $3.8e - 20$ |
| WM_pve | $0.0654 \pm 0.1719$ | $0.0144 \pm 0.0277$ | *ns* | $0.9675 \pm 0.0158$ | $0.8634 \pm 0.0870$ | $1.3e - 17$ |
| WM_bin | $0.0952 \pm 0.1669$ | $0.0096 \pm 0.0086$ | *ns* | $0.9530 \pm 0.0193$ | $0.8322 \pm 0.0822$ | $9.0e - 19$ |
| WM_Skel_pve | $0.0803 \pm 0.0899$ | $0.1571 \pm 0.1569$ | *ns* | $0.9132 \pm 0.0262$ | $0.8654 \pm 0.0229$ | $2.4e - 15$ |
| WM_Skel_bin | $0.0495 \pm 0.0641$ | $0.0677 \pm 0.0907$ | *ns* | $0.8777 \pm 0.0353$ | $0.8129 \pm 0.0254$ | $3.3e - 16$ |
| **T2 high** |  |  |  |  |  |  |
| GM_pve | $0.0042 \pm 0.0042$ | $0.0042 \pm 0.0038$ | *ns* | $0.9190 \pm 0.0287$ | $0.8548 \pm 0.0634$ | $7.2e - 09$ |
| GM_bin | $0.0069 \pm 0.0077$ | $0.0106 \pm 0.0131$ | *ns* | $0.9034 \pm 0.0314$ | $0.8395 \pm 0.0578$ | $9.0e - 10$ |
| GM_Skel_pve | $0.0305 \pm 0.0523$ | $0.0336 \pm 0.0379$ | *ns* | $0.8767 \pm 0.0272$ | $0.8344 \pm 0.0185$ | $1.7e - 16$ |
| GM_Skel_bin | $0.0230 \pm 0.0395$ | $0.0379 \pm 0.0340$ | 0.0005 | $0.8471 \pm 0.0332$ | $0.8119 \pm 0.0220$ | $9.0e - 10$ |
| WM_pve | $0.0074 \pm 0.0063$ | $0.0052 \pm 0.0046$ | *ns* | $0.9158 \pm 0.0274$ | $0.8508 \pm 0.0607$ | $6.6e - 10$ |
| WM_bin | $0.0122 \pm 0.0140$ | $0.0140 \pm 0.0217$ | *ns* | $0.8850 \pm 0.0328$ | $0.8247 \pm 0.0539$ | $8.2e - 10$ |
| WM_Skel_pve | $0.0166 \pm 0.0230$ | $0.0380 \pm 0.0577$ | *ns* | $0.8664 \pm 0.0364$ | $0.8317 \pm 0.0189$ | $7.3e - 10$ |
| WM_Skel_bin | $0.0388 \pm 0.0543$ | $0.0149 \pm 0.0219$ | *ns* | $0.8318 \pm 0.0437$ | $0.7923 \pm 0.0249$ | $7.3e - 09$ |
| **T2 low** |  |  |  |  |  |  |
| GM_pve | $0.1076 \pm 0.1886$ | $0.0071 \pm 0.0060$ | *ns* | $0.9574 \pm 0.0238$ | $0.8713 \pm 0.0884$ | $6.8e - 13$ |
| GM_bin | $0.0690 \pm 0.1343$ | $0.0163 \pm 0.0182$ | *ns* | $0.9388 \pm 0.0294$ | $0.8434 \pm 0.0843$ | $3.3e - 14$ |
| GM_Skel_pve | $0.0179 \pm 0.0192$ | $0.0519 \pm 0.0843$ | *ns* | $0.8423 \pm 0.0429$ | $0.7795 \pm 0.0364$ | $1.7e - 12$ |
| GM_Skel_bin | $0.0163 \pm 0.0202$ | $0.0389 \pm 0.0586$ | *ns* | $0.8177 \pm 0.0456$ | $0.7465 \pm 0.0368$ | $9.5e - 14$ |
| WM_pve | $0.0816 \pm 0.1741$ | $0.0095 \pm 0.0126$ | *ns* | $0.9405 \pm 0.0267$ | $0.8456 \pm 0.0905$ | $2.7e - 13$ |
| WM_bin | $0.0614 \pm 0.1084$ | $0.0096 \pm 0.0112$ | *ns* | $0.9176 \pm 0.0358$ | $0.8107 \pm 0.0864$ | $2.4e - 14$ |
| WM_Skel_pve | $0.0369 \pm 0.0374$ | $0.0465 \pm 0.0735$ | *ns* | $0.8108 \pm 0.0574$ | $0.7296 \pm 0.0376$ | $3.3e - 14$ |
| WM_Skel_bin | $0.0359 \pm 0.0649$ | $0.0131 \pm 0.0243$ | *ns* | $0.8137 \pm 0.0526$ | $0.7212 \pm 0.0479$ | $1.4e - 14$ |

**Figure 3:**
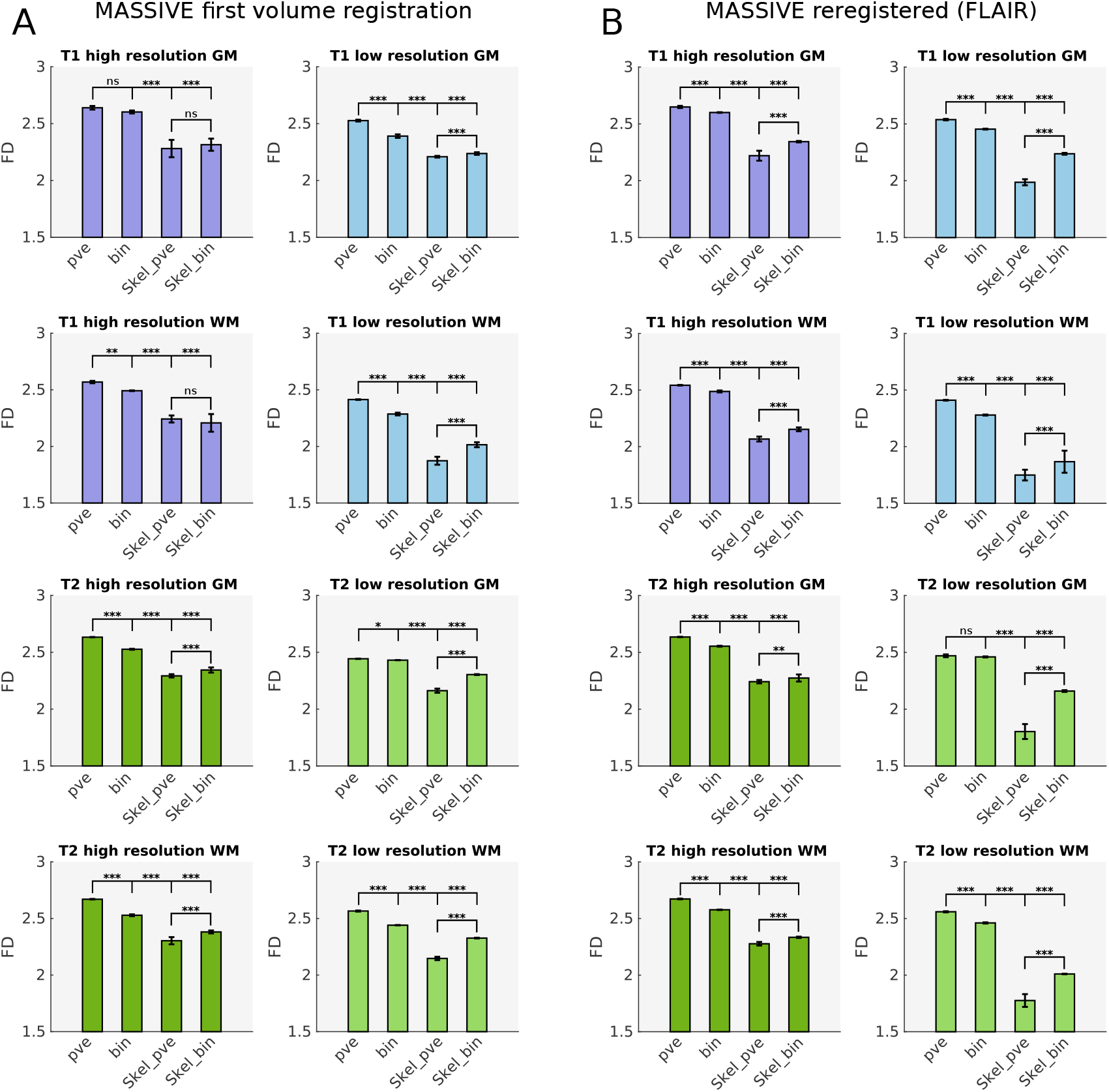
Parameter-dependent comparison of the fractal dimension estimates in the MAS-SIVE data set. Panels A and B summarize the comparisons of the mean fractal dimension estimates over image analysis groups in first volume and FLAIR registration, respectively. Horizontal bars reflect pair-wise significance levels. Comparisons for binary-segmented images invariably yielded the same significance levels as the partial volume estimates so they were omitted here for visual coherence, as in fig. 12. Note the remarkably similar effects regarding binarization, tissue type, and skeletonization as compared to the results from the MSC data in the main text. *ns*: not significant; *: *p <* 0.05; **: *p <* 0.01; ***: *p <* 0.001; bin: binary tissue segmentation; GM: gray matter; pve: partial volume estimates; Skel: skeleton model; WM: white matter.

**Figure 4:**
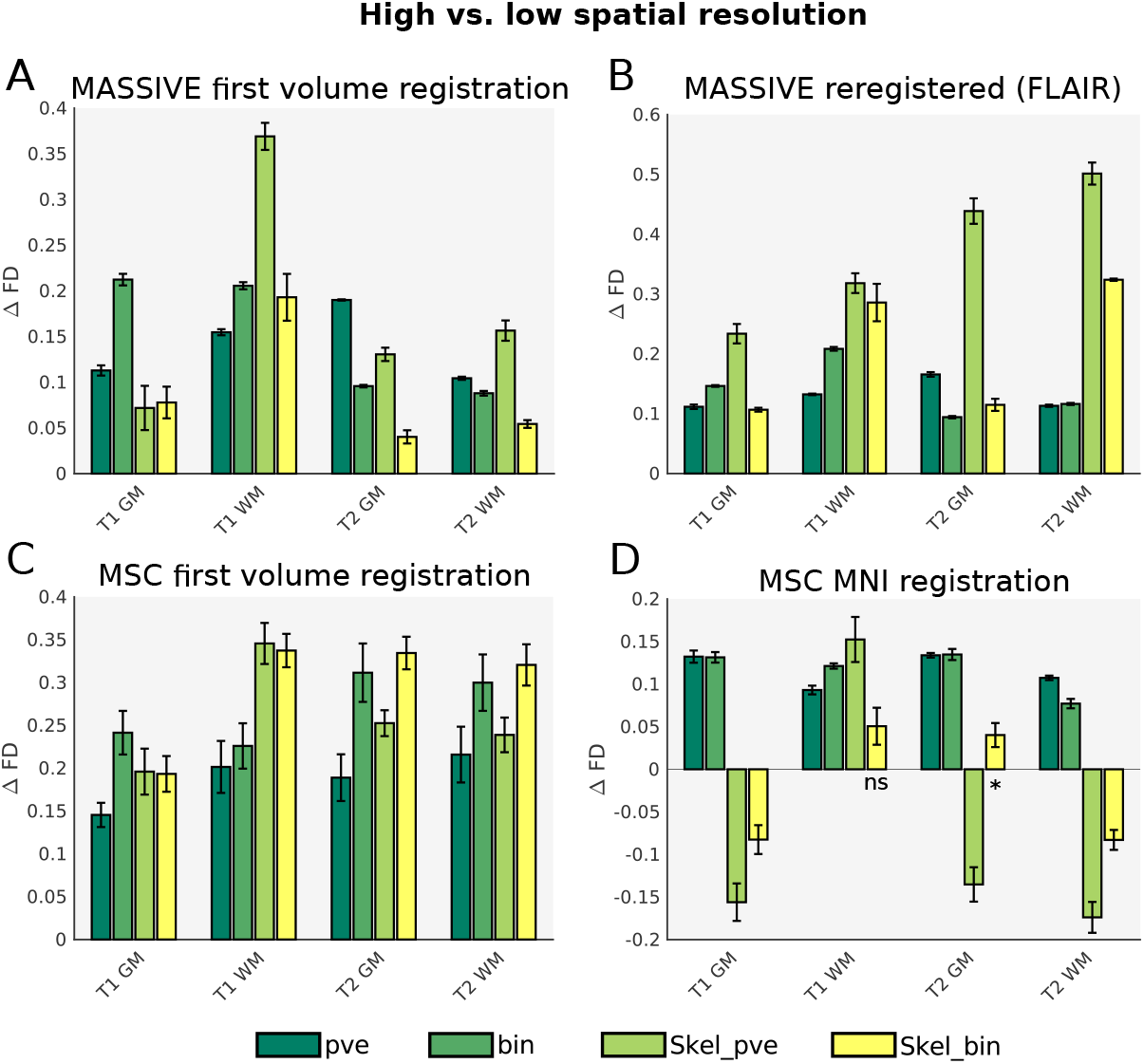
Fractal dimension differences in high vs. low spatial resolution. Bars represent the respective difference in the mean fractal dimensions of high and low resolution images for the MASSIVE (panels A and B) and the MSC (panels C and D) data sets in both image registrations. Error bars correspond to the difference’s sampling distribution error. All high vs. low resolution comparisons were significant at *p <* 0.001 unless otherwise indicated (*: *p <* 0.05; *ns*: not significant.). For unskeletonized images, the fractal dimensions were significantly decreased by lower spatial resolution in both data sets, regardless of image registration or tissue type. The same was true for most comparisons concerning skeleton models with a few exception in the MSC data in MNI registration (panel D). bin: binary tissue segmentation; ∆*FD*: difference in mean fractal dimension in high vs. low spatial resolution; GM: gray matter; pve: partial volume estimates; Skel: skeleton model; WM: white matter.

**Figure 5:**
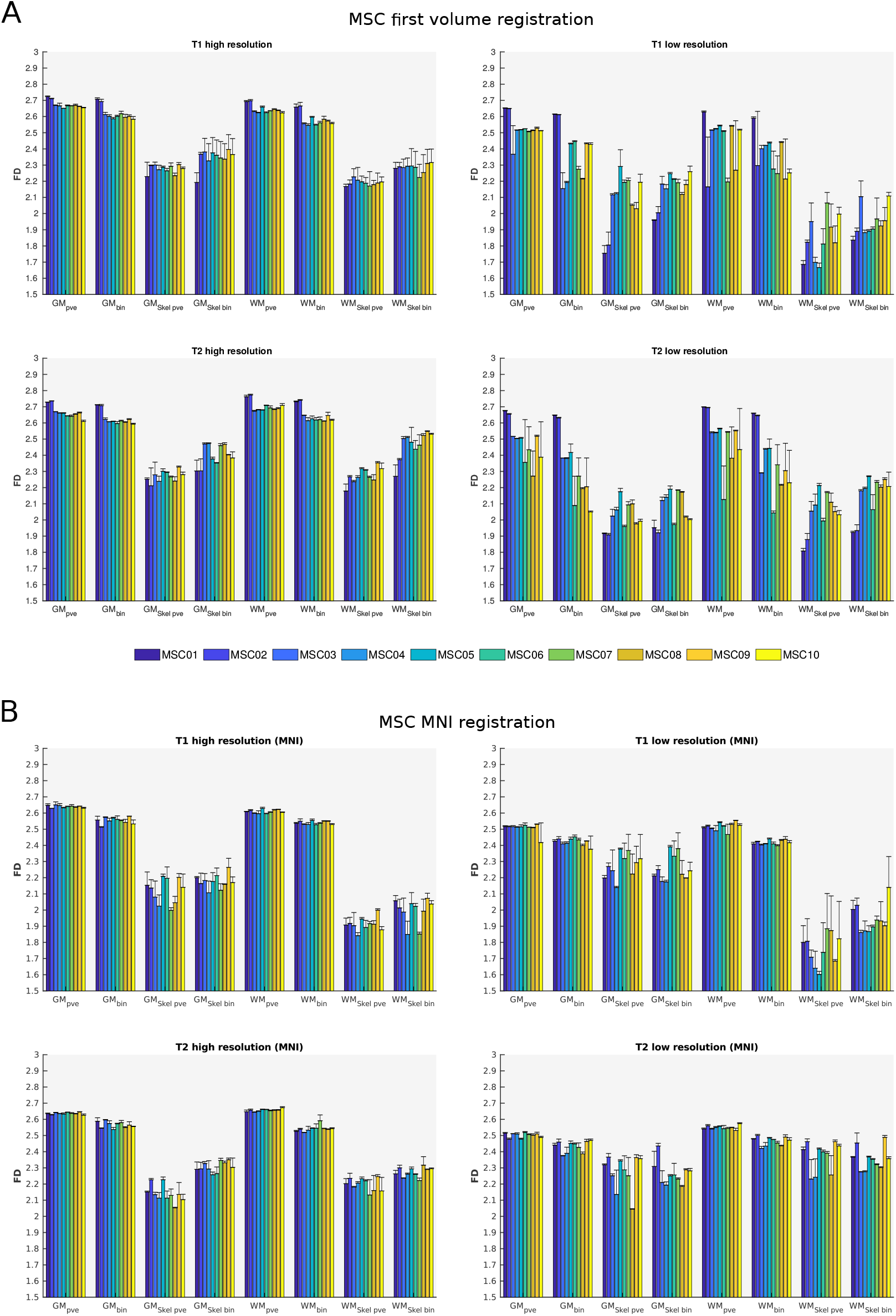
Results of the fractal dimension estimation in the MSC data set. The figure visualizes the fractal estimation results across image registration (panel A: first volume registration; panel B: MNI registration), subjects (grouped bars) and analysis groups. Note the higher between-subject variability in first volume as compared to MNI registration and the higher within- and between-subject variability in skeleton models as opposed to unskeletonized images. GM: gray matter; bin: binary tissue segmentation; pve: partial volume estimates; Skel: skeleton model; WM: white matter.

